# Mineralocorticoid and glucocorticoid receptor interaction drives TGFβ1-induced triple-negative breast cancer progression to metastasis

**DOI:** 10.1101/2025.11.03.686330

**Authors:** Sai Harshita Posani, Caroline H. Diep, Raisa I. Krutilina, Hilaire Smith, Tiffany N. Seagroves, John Blenis, Carol A. Lange

**Affiliations:** Graduate program in Pharmacology, University of Minnesota, Minneapolis, MN 55455 USA; Masonic Cancer Center, University of Minnesota, Minneapolis, MN 55455, USA; Tulane University School of Medicine, Department of Medicine, New Orleans, LA, USA 70112; Tulane Cancer Center, Tulane University, School of Medicine, New Orleans, LA, USA 70112; Department of Pharmacology, Weill Cornell Medicine, New York, NY; Department of Medicine, Division of Hematology, Oncology & Transplantation, University of Minnesota, Minneapolis, MN 55455, USA

**Keywords:** triple negative breast cancer, mineralocorticoid receptor (NR3C2 gene), glucocorticoid receptor (NR3C1 gene), TGFβ1, finerenone

## Abstract

In triple-negative breast cancer (TNBC), p38 MAPK phosphorylates the glucocorticoid receptor (GR) at N-terminal Ser134 in response to cytokines, such as TGFβ1. Phospho-Ser134-GR (pSer134-GR) regulates genes that promote migratory/invasive behavior and altered metabolism. In addition to acting as ligands for GR, glucocorticoids also activate closely-related mineralocorticoid receptors (MR). Elevated MR activity via its physiological ligand aldosterone (aldo), mediates hypertension, inflammation and fibrosis. While GR/p-GR has been implicated in TNBC progression to metastasis, the contribution of MR remains unknown. The METABRIC dataset demonstrated significantly higher expression of MR transcripts in TNBC relative to luminal breast cancers. Further, high MR expression predicted worse overall survival in the KM-plotter database. We observed pSer134-GR- and p38-dependent association of cytoplasmic MR/p-GR complexes upon treatment of TNBC cells with TGFβ1, while nuclear MR-GR complexes predominated in response to dexamethasone (dex) and/or aldo. Cytoplasmic MR/p-GR complexes entered the nucleus within 4 hours. MR knockdown or inhibition with MR-selective antagonists (spironolactone, finerenone) significantly reduced aldo or TGFβ1-induced migratory and stemness properties. MR knockdown models exhibited reduced migration, attenuated stem cell expansion, and impaired metastasis to lungs following tail-vein injection, thereby phenocopying cells harboring phospho-mutant S134A-GR. MR activation by aldo or TGFβ1 transcriptionally upregulated both canonical MR-target genes (SGK1, ENaC1) and non-canonical target genes related to fibrosis (COL1A1, ICAM1 and CTGF). MR expression was essential for functionally intact p38 MAPK/p-Ser134-GR signaling downstream of TGFβ1 receptor activation. Our studies define a novel role of MR/p-GR complexes in regulation of TNBC cell migration, stemness potential, and metastasis. Pharmacological inhibition of MR with FDA-approved MR antagonists (MRAs) offers an exciting opportunity for improved clinical management of TNBC patients.

## Introduction

Cancer remains a global health concern due to its rising incidence and profound impact on physical, mental, and social well-being (1). Breast cancer (BC) is the most prevalent malignancy in women, with a 12.5% incidence rate in the United States (2). BC is a molecularly and clinically heterogeneous disease, and current treatments are guided by the presence or absence of estrogen receptors (ER), progesterone receptors (PR) and elevated HER2 receptor expression characterized by HER2 gene amplification (3). Gene expression profiling has classified BC into five subtypes: luminal A, luminal B, HER2-enriched, basal-like, and normal-like (4, 5). Basal-like tumors typically exhibit a triple-negative phenotype (ER−, PR−, HER2−), known as triple-negative breast cancer (TNBC), accounting for approximately 15% of all breast cancers (6). TNBC is associated with aggressive behavior, a high risk of distant metastasis, and poor survival outcomes, with a median overall survival of 12 months and a four-year survival rate below 20% (7). Current TNBC treatments include cytotoxic chemotherapy, immunotherapies, PARP inhibitors, and antibody-drug conjugates such as sacituzumab govitecan, which targets the TROP2 cell surface receptor conjugated to a topoisomerase inhibitor payload (8, 9). While these therapies can be highly beneficial in selected TNBC patients, effective targeted therapies remain limited. As a result, TNBC patients experience higher recurrence and metastasis rates, and worse prognosis compared to other subtypes (10).

Glucocorticoid receptors (GR) are steroid hormone receptors that are overexpressed in endocrine-related cancers such as ovarian, prostate, pancreatic and breast cancer (11). High expression of GR is correlated with a worse prognosis in TNBC, but with a better prognosis in early-stage ERα-positive breast cancers (12, 13). A meta-analysis of 1,378 early-stage ERα-negative and 623 TNBC cases showed that high GR expression was significantly associated with poor relapse-free survival, irrespective of chemotherapy status (14). As with other steroid receptors (SRs), GR activity is modulated by post-translational modifications (15). We have demonstrated that phosphorylation of GR at an N-terminal serine 134 (Ser134) is elevated in TNBC compared to other subtypes (16). We have also shown that cellular stress signals relevant to cancer progression (hypoxia, nutrient starvation, local pro-inflammatory factors including cytokines (TGFβ1), cytotoxic chemotherapies), activate the p38 MAPK pathway, thereby driving persistent GR Ser134 phosphorylation as a required event preceding TNBC cell migration, invasion and secondary tumorsphere formation, an indirect measure of BC cell stemness properties (16). Importantly, this study identified MAP3K5, also known as mitogen-activated protein kinase kinase kinase 5, MAPK/ERK kinase kinase 5, MEKK5, or apoptosis signal-regulating kinase 1, ASK1 as a key downstream target gene regulated by p-Ser134-GR. Expression/activity of MAP3K5 was required for robust activation of the downstream MEK3/6-p38 MAPK signaling module. Taken together, these landmark studies placed p-Ser134-GR as a key effector of TGFβ1 signaling in TNBC cells.

In addition to binding GR, corticosteroids are agonists for the mineralocorticoid receptor (MR) (17). MR is a well-studied member of the steroid receptor superfamily of ligand-activated transcription factors that primarily functions to control sodium and potassium transport in epithelial cells, more specifically in the kidney and colon (18). The principal endogenous ligand for MR is aldosterone (aldo), which regulates fluid and electrolyte balance in kidneys, salivary glands, sweat glands and colon (19). Aldo, the end-product of the renin-angiotensin-aldosterone system (RAAS) in the zona glomerulosa of the adrenal cortex, regulates blood pressure and fluid/electrolyte balance by activating MR, thereby controlling sodium reabsorption and potassium secretion (20). In mouse models of chronic blood pressure overload and myocardial infarction, deletion or inactivation of the MR gene attenuated left ventricular dilatation, cardiac hypertrophy, and development of heart failure. Overexpression of MR in cardiomyocytes induced ventricular remodeling, development of fibrosis, heart failure and pro-arrhythmogenic effects (21–23). In two randomized clinical trials, MR blockade significantly demonstrated reduced morbidity and mortality in heart failure patients (24, 25). Overexpressing MR in cardiomyocytes accelerates adverse cardiac remodeling, heart failure, and arrhythmias (26). Excess aldo, via hyperactivation of MR, underlies numerous cardiovascular, adipose, and renal disease states, including inflammation, fibrosis, and vascular dysfunction (27, 28). Moreover, in vascular smooth muscle cells, non-canonical MR signaling leads to vasoconstriction via phosphorylation of ERK, increased intracellular calcium levels, and enhanced production of reactive oxygen species via activation of IGF-1, c-SRC, p38 MAP kinase and PI3K signaling (29, 30). Crosstalk with the stress-activated receptors appears to be a common theme for non-canonical cardiovascular effects of MR (31, 32).

MR antagonists have been foundational in clinical practice for cardiovascular and renal indications and are now being explored for broader therapeutic applications, such as for treatment of fibrosis (33, 34). Spironolactone (spiro), a first-generation steroidal MR antagonist, exhibits potent MR blockade, but the use of this agent is limited by off-target antiandrogenic and progestogenic effects, leading to gynecomastia and menstrual irregularities (35, 36). Eplerenone, a more selective steroidal antagonist, retains efficacy in heart failure while displaying a reduced side-effect profile (37). More recently, non-steroidal MR antagonists such as finerenone (fine) have demonstrated superior tissue selectivity with a lower risk of hyperkalemia, earning regulatory approval for reducing the progression of diabetic nephropathy and decreasing cardiovascular morbidity in patients with type 2 diabetes and chronic kidney disease (38, 39).

Recent exciting studies have shed light on the mechanisms and consequences of GR-MR interactions in physiology. GR and MR can interact to form heterodimers, influencing gene expression in a context-dependent manner (17, 40). The heterodimers exhibit distinct transcriptional activities compared to their homodimer counterparts, affecting genes involved in stress responses and homeostasis (41). The formation and function of GR-MR heterodimers are influenced by ligand availability and cellular context. For instance, in keratinocytes, GR-MR heterodimers synergistically activate specific gene promoters in response to the GR ligand dexamethasone (dex) (42). Similarly, in the hippocampus and renal cells, these receptors co-occupy gene regulatory promoters such as PER1, leading to unique transcriptional outcomes (43). Unstimulated MR exists as a monomer and forms a tetramer upon aldo stimulation (44), whereas to form MR and GR heterocomplexes, GR displaces MR subunits (44). Such tissue-specific interactions underscore the complexity of GR-MR crosstalk in regulating physiological processes. In multiple myeloma, crosstalk between GR and MR enhances the efficacy of glucocorticoid-induced apoptosis (45). Specifically, dex downregulates MR levels, and co-treatment with the MR antagonist spiro amplifies cell death in glucocorticoid-sensitive myeloma cells. The role of MR in TNBC is entirely unknown.

Herein, we sought to evaluate the requirement for MR in promoting aggressive *in vitro* breast cancer phenotypes in GR+/MR+ TNBC models, such as increased cell migration, formation of secondary tumorspheres indicative of stemness properties, and lung colonization following mouse tail vein injection of human TNBC cells *in vivo*. Biological responses were compared in response to pharmacological inhibition and genetic knockdown of MR using shRNA. We also queried the regulation of MR-GR close-proximity interactions in the context of hormone and cytokine (TGFβ1) treatments and p38 MAPK-dependent phosphorylation of GR-Ser134 owing to the significant attention that MR and GR crosstalk has garnered due to their overlapping roles in gene regulation and implications for related physiological and pathological processes in health and disease.

## Methods

### RNA expression analysis for METABRIC dataset samples

Molecular Taxonomy of Breast Cancer International Consortium (METABRIC) (via cBioPortal – http://www.cbioportal.org/), data were obtained for patients who had the 3-gene classifier and microarray data available for GR and MR. mRNA expression values were log2 transformed and then boxplots were created and analyzed using GraphPad Prism to compare expression levels across groups. Statistical significance was calculated based on one-way ANOVA, followed by Tukey post-hoc testing.

### Survival analysis from KM-plotter

The KM plotter database was used to evaluate survival analysis - https://kmplot.com/analysis (46). Patients with estrogen, progesterone and HER-2 receptor-negative tumors for whom overall survival data were reported were selected for analysis based upon MR expression (n=126) and were stratified into two groups based on the “auto best cut-off setting.”

### Cell lines and culture conditions

Parental (unmodified) cells were obtained from Dr. Tiffany Seagroves and maintained in the indicated base medium supplemented with 10% FBS (Corning, cat# 35-010-CV), and 1% penicillin-streptomycin (Gibco, cat# 15140-122): MDA-MB-231 (DMEM-Hi Corning cat# MT10013CV), SUM159 (DMEM F/12 Corning cat# MT15090CV, 1 mg/mL insulin Sigma Aldrich cat# I6634, 1 mg/mL hydrocortisone MedChemExpress cat# HY-N0583), Hs578T (DMEM-Hi Corning cat# MT10013CV, 10 mg/mL insulin Sigma Aldrich cat# I6634) and MDA-MB-468 cells (DMEM/F12 Corning cat# MT15090CV). All cell lines were authenticated by the University of Arizona Genomics Core (RRID: SCR_012429), and were tested negative for mycoplasma (Lonza, MycoAlert assay, cat# LT07-418). For experiments requiring hormone treatment, cells were starved for 18-20 hours in iMEM (Gibco, cat# A1048801) supplemented with 1% dextran-coated charcoal FBS (DCC) (HyClone, cat# SH3006803). All treatment conditions are included in the figure legends.

### Western blotting

At experimental endpoints, the media was removed, and cells were washed thrice with 1× PBS. Whole cell lysates were isolated using a radioimmunoprecipitation (RIPA) assay buffer (RIPA lite) supplemented with 1 mmol/L PMSF, 5 mmol/L NaF, 0.05 mmol/L Na3VO4, 25 mmol/L ß-glycerophosphate (BGP), 20 μg/mL aprotinin, 1 complete mini tablet protease inhibitors (Roche, cat# 11836153001) and 1 tablet of PhosSTOP (Roche cat# 4906837001). Protein quantified by Bradford assay was loaded (30 µg per lane) and resolved by 10% SDS-PAGE gels, followed by transfer to polyvinylidene difluoride membranes. Blots were probed with the following primary antibodies: custom pGR (1:750, Pierce Biotechnology), GR (1:1,000, Santa Cruz Biotechnology, cat# sc-393232), MR (1:1,000, Cell Signaling technology cat# 58883S), p-p38 (1:1,000, Cell Signaling Technology cat #4511S), p38 (1:1,000, Cell Signaling Technology cat# 9212), and GAPDH (1:5,000, Santa Cruz Biotechnology, cat# sc-0411). Horseradish peroxidase– conjugated goat anti-rabbit (Bio-Rad cat# 170-6515) or goat anti-mouse (Bio-Rad cat# 170-6516,) were used as secondary antibodies. All blots were exposed to ECL reagent (SuperSignal West Pico PLUS; Thermo Fisher Scientific cat# PI34580) followed by detection by film.

### Proximity ligation assay (PLA)

Proximity ligation assay was performed using the Duolink™ In Situ Red Kit Mouse/Rabbit (Millipore Sigma, cat# DUO92008-100RXN). MDA-MB-231 cells were plated on glass coverslips in 6-well plates. After the indicated endpoints, the slides were washed thrice with cold 1x PBS, fixed with 4% paraformaldehyde for 15 min at RT and then permeabilized (0.1% Triton X-100, 1×PBS) for 15 min at RT. Slides were then incubated with the Duolink blocking solution for 1 hour at 37°C, followed by incubation with primary antibodies (MR – 1:500; GR - 1:250) overnight at 4 °C, incubation with secondary PLA probes (anti-rabbit PLA-minus and anti-mouse PLA-plus) and washing. The ligase and amplification steps were performed according to the manufacturer’s protocol. Samples were then washed and mounted with the Duolink® In Situ Mounting Medium with DAPI (Millipore Sigma, cat# DUO82040-5ML). For each treatment condition, a negative control was performed, omitting the anti-MR antibody. To quantify the PLA signals (as puncta), images were acquired using a Keyence microscope and analyzed using Fiji ImageJ. The same size threshold for the nuclei and PLA puncta was applied equally to all images. For each region investigated, the number of puncta measured was divided by the number of nuclei in the area.

### Cell migration

Cell migration was assessed by 8 µm transwell inserts (Corning cat# 353097). Following starving, cells were trypsinized and plated in iMEM media at a density of 5 x 10^4^ cells into the upper chamber, and 10% FBS was added as the chemoattractant agent in the lower well. All treatments were added in the presence of 1% DCC as indicated in the figure legends. MDA-MB-231 and SUM159 cells were allowed to migrate for either 6 or 18 hours before fixation with 4% paraformaldehyde and staining with hematoxylin (Sigma Aldrich cat# HHS16). The number of migrated cells was counted over 4 representative fields per sample at 10X magnification on a light microscope, and all results were normalized to the number of migrated vehicle cells.

### Tumorsphere assay

Cultured cells were digested to single cells using 0.25% trypsin-EDTA and then strained through a 40 μm sieve. Cells were plated in ultra-low attachment plates (Corning cat# 3473) at 1 × 10^3^ cells per well and cultured in defined phenol red-free DMEM/F12 medium (Corning, cat# 16-405-CV) containing 1% methylcellulose (Sigma Aldrich cat# M7207), 1% charcoal-stripped B27 supplement (Invitrogen), 1% penicillin-streptomycin, 20 ng/mL EGF (Sigma Aldrich), 20 ng/mL basic-FGF (Gibco), and 10 μg/mL heparin (Sigma Aldrich). After 5–7 days, primary spheres were quantified, harvested and digested into single cells to generate secondary tumorspheres as described previously (47). The secondary tumorspheres were allowed to grow for 5–7 days before manual counting. Data are presented as the average ± SD of three biological replicates.

### Quantitative RT-PCR

Quantitative real-time PCR (RT-qPCR) experiments were conducted as previously described (16). Following treatments, the media was removed, and cells were washed thrice with cold 1× PBS. RNA was isolated with Trizol, as per the manufacturer’s protocol (Roche cat# 11667165001). cDNA was generated from total RNA extracted from MDA-MB-231 and SUM159 cells expressing either shGFP or shMR constructs using qScript cDNA SuperMix (Quanta Biosciences cat# 101414-108). RT-qPCR was performed using Light Cycler FastStart DNA Master SYBR Green I (Roche cat# 06924204001) on a LightCycler 96 Real-Time PCR system (Roche). The cycling conditions by qPCR were as follows: 10 min of initial denaturation at 95°C, 10 sec denaturation at 95°C, 10 sec at 60°C for annealing of primers and extension at 72°C for 5 sec for 45 cycles. Relative target gene expression was normalized to the expression of the internal control gene 18S mRNA. Primer sequences are indicated in Supplementary Figure 1A. Data are presented as the average ± SD of three independent measurements.

### Generation of MDA-MB-231 shGFP and shMR stable knockdown cells

Stable shMR cells (TRCN0000019364 and TRCN0000019366) were generated by transducing MDA-MB-231 models with CMV-Neo lentiviral vectors containing target gene shRNA sequences (MISSION TRC library, Sigma). Transduced cells were selected and maintained 1 μg/mL puromycin (Sigma Aldrich cat# P8833).

### Generation of siRNA mediated MR knockdown in MDA-MB-231 cells

To transiently deplete MR, MDA-MB-231 cells were plated at a density of 2x10^5^ cells per well in 6-well plates and grown to 80% confluence. Cells were then transfected with pre-incubated mixtures of either control siRNA (cat# EHU088591-50UG) or siRNA to MR (6.8 mL of siRNA/well plus 7.5 mL/well of transitX2 transfection reagent) (Eupheria Biotech cat# 10766-892) in 250 mL/well of OptiMEM medium (lnvitrogen) and allowed to incubate for 24 hours at 37°C. Whole cell lysates and RNA were then collected as described previously to evaluate MR protein and RNA levels.

### Experimental metastasis (tail vein) assays

All animal studies are approved by the Tulane University local IACUC (protocol #2186) and are consistent with US Public Health Service (PHS) Policy and the Animal Welfare Act (AWA). Female Nod/Scid/Gamma mice (Jax Labs, stock #005557; RRID:IMSR_JAX:005557) fed Teklad 7904 breeder diet (Inotiv) ad libitum were injected in the tail vein with 250,000 shGFP or shMR cells at 8-9 weeks of age. Prior to injection, cells were stained with diR lipophilic dye (Biotium cat #60017) in order to confirm successful injections (signal is restricted to the lungs) using an IVIS Lumina III bio-imager. Only mice with signal in lungs >1 x 10^9^ total photon flux were kept on study (n=7 shGFP and n=7 shMR). Animals were euthanized 21 days later from anesthetized animals. The lungs were inflated with saline prior to removal, followed by histopathology at the Tulane Department of Pathology Histology Laboratory and FFPE lung sections were stained with H&E. Whole H&E-stained slides were digitized at the University of Minnesota University Imaging Core on a Huron TissueScope LE instrument at 20x magnification, followed by quantification of metastatic burden in whole lung sections using QuPath open-source software (v 0.6.0; RRID:SCR_018257) (48).

### General Reagents

Aldosterone (aldo; Cayman Chemical, cat# 15273), dexamethasone (dex; Sigma Aldrich; cat# D4902), spironolactone (spiro; Cayman Chemical, cat# 9000324), and hydrocortisone (Sigma-Aldrich) stocks were prepared in ethanol. Finerenone (fine; Cayman Chemical cat# 36086) stocks were prepared in sterile dimethyl sulfoxide (Sigma-Aldrich cat# D8418). Epidermal growth factor (EGF; Sigma-Aldrich, cat# E9644) was prepared in 0.1% bovine serum albumin containing 10 mM acetic acid. Human FGF basic 146 aa recombinant protein (FGF2; R&D Systems, cat# 233FB025CF) was prepared in sterile PBS. TGFβ1 (R&D Systems cat# 7754-BH) was prepared in sterile PBS.

### Statistics

All statistical approaches are disclosed in the legends after analysis using Prism (GraphPad, v7-8). For experiments that involved only two groups, two-tailed Student’s *t*-tests were performed, and when more than two groups were compared, one-way or two-way ANOVA testing was utilized, followed by post-hoc Dunnett’s or Tukey tests to identify which comparisons among the groups were significant.

## Results

### Expression of MR is elevated in triple-negative breast cancer (TNBC) public datasets and TNBC cell lines

To investigate the role of mineralocorticoid receptor (MR; gene name: NR3C2) in TNBC, we analyzed mRNA expression across breast cancer subtypes using publicly available datasets. Expression data from the METABRIC dataset were retrieved via cBioPortal. MR mRNA levels were elevated in patients with ER−/HER2− and HER2+ breast cancers compared to those with ER+/HER2− tumors (Figure 1A). Previous studies have reported that high cytoplasmic MR expression correlates with poor relapse-free survival (RFS) in ER+ breast cancer patients (49). Additional research has linked MR expression to longer OS in non-small cell lung and colorectal cancers (50, 51). We used the KM-plotter tool to assess the relationship between MR mRNA expression and survival outcomes in TNBC cohorts. The “auto select best cutoff value” option was chosen to stratify patients into two comparison groups, which searches for the optimal cutoff values between lower and upper quartiles. This analysis revealed that patients with ER−/PR−/HER2− BC expressing high levels of MR mRNA (n = 78) had significantly worse OS than patients with lower levels of MR (n = 48) (Figure 1B).

**Figure 1:**
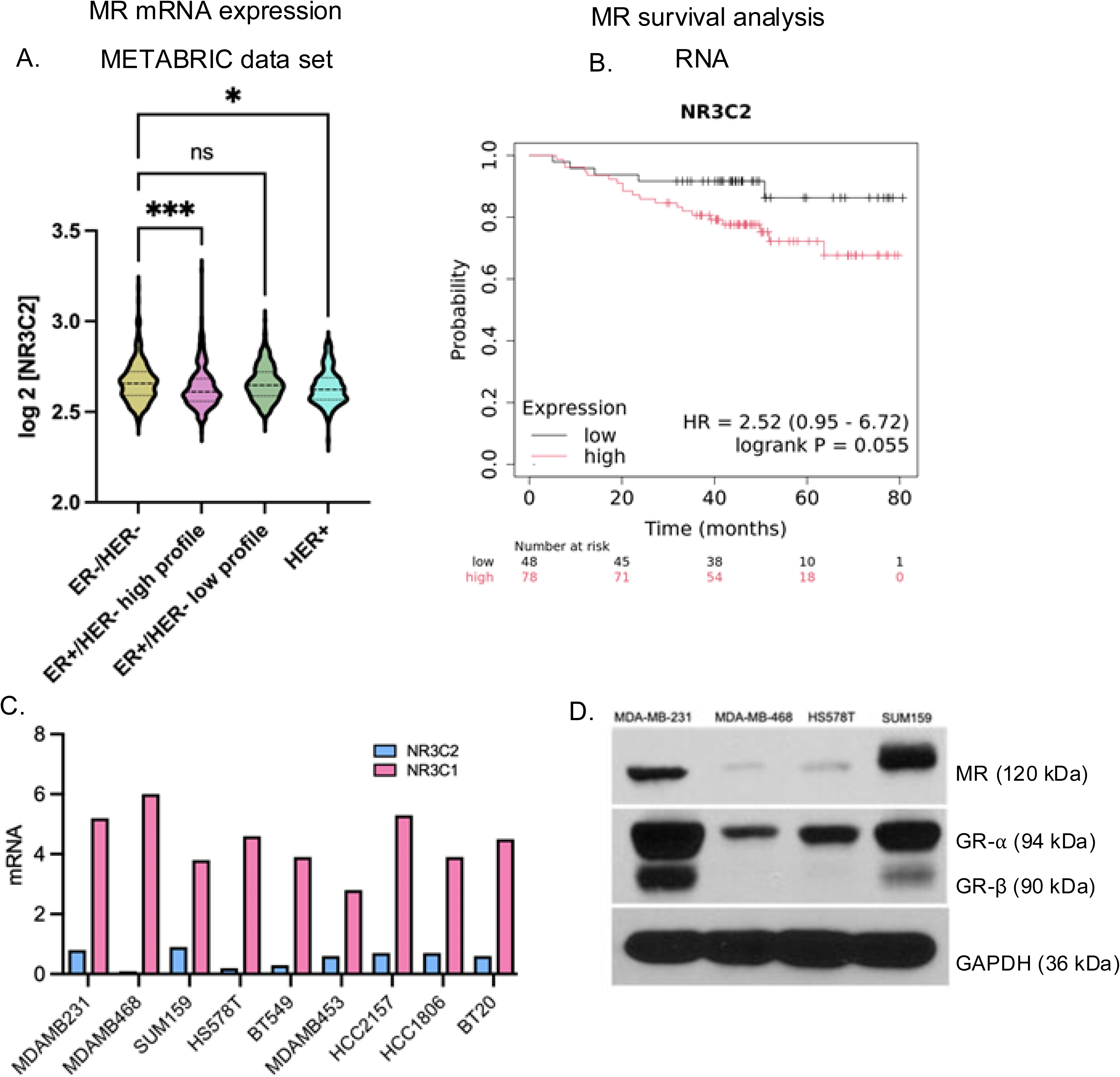
(A) mRNA Expression of MR (*NR3C2*) in breast cancer patients was analyzed using the METABRIC database (n=1,700). Significance was assessed by one-way ANOVA and Tukey testing (*P < 0.05, **P < 0.01, ***P < 0.001, ****P < 0.0001). (B) Overall survival analysis for patients from the KM-plotter database using RNA seq data. Patients were stratified into two groups based on the best-fit algorithm to stratify the patients into two groups (“high expression”, n=78) or (“lower expression”, n=48) and analyzed for significance by log-rank testing (*P < 0.05, **P < 0.01, ***P < 0.001, ****P < 0.0001). (C) mRNA expression of MR (*NR3C2*) and GR (*NR3C1*) in TNBC cell lines from the cancer cell line encyclopedia (CCLE). (D) Representative Western blot analysis of MR, GR-α and GR-β in selected TNBC cell lines.

To evaluate the expression of MR (NR3C2) and GR (NR3C1) across multiple TNBC cell line models, we analyzed transcriptomic data from the Cancer Cell Line Encyclopedia (CCLE). NR3C1 (GR) mRNA was expressed at moderate to high levels in most TNBC lines, whereas NR3C2 (MR) mRNA expression was moderate to low (Figure 1C). We selected those cell lines that expressed high levels of GR and moderate levels of MR for further study. We next assessed basal MR and GR protein expression in a panel of four TNBC cell lines by immunoblotting (Figure 1D). Densitometry values are shown in Supplementary Figure 2A and 2B MDA-MB-231 and SUM159 cells expressed high to moderate basal levels of at least two GR isoforms (GR-α and GR-β) and MR protein whereas MDA-MB-468 and Hs578T cells expressed low levels of MR and only the GR-α isoform. GR-α and GR-β are produced by alternative splicing of exon 9 of NR3C1. They share the same first 727 amino acids but diverge at their C-terminal ends (52).

### MR and GR associate in close-proximity upon steroid hormone or TGFβ1 treatment

Although traditionally studied as independent pathways, increasing evidence suggests that MR and GR can physically interact, forming heterodimers or higher order complexes that modulate transcriptional activity (17). We employed PLA to investigate MR-GR interactions in TNBC across different cellular conditions to elucidate how their association is regulated. In vehicle (veh) treated MDA-MB-231 cells, few to no MR-GR PLA signals (puncta) appeared, whereas following either 1 µM dexamethasone (dex) or 1 µM aldosterone (aldo) treatment for 90 min, red puncta indicating MR-GR interactions appeared that were predominantly localized to the nucleus (Figure 2A). When 1 µM dex and 1 µM aldo were added simultaneously, MR-GR interactions dramatically increased relative to single ligands (Figure 2A). In SUM159 cells, PLA signals appeared in the nucleus upon either 1 µM dex or 1 µM aldo treatment and signals increased significantly with the combination of both (Figure 2B). Interestingly, in stark contrast to steroid hormone treatment, 10 ng/mL TGFβ1 treatment induced an increased number of MR-GR puncta that were localized to the cytoplasm (Figure 2A and 2B). This suggests that MR-GR interactions are initiated or stabilized in the cytoplasm in response to TGFβ1, possibly as part of a pre-translocation complex. Alternatively, MR-GR complexes may function as mediators in cytoplasmic signaling independently of their classical nuclear transcriptional functions. This cytoplasmic retention and interaction imply a non-genomic rapid signaling function for MR-GR complexes, consistent with increasing recognition of essential extranuclear roles for other steroid receptor family members (53).

**Figure 2:**
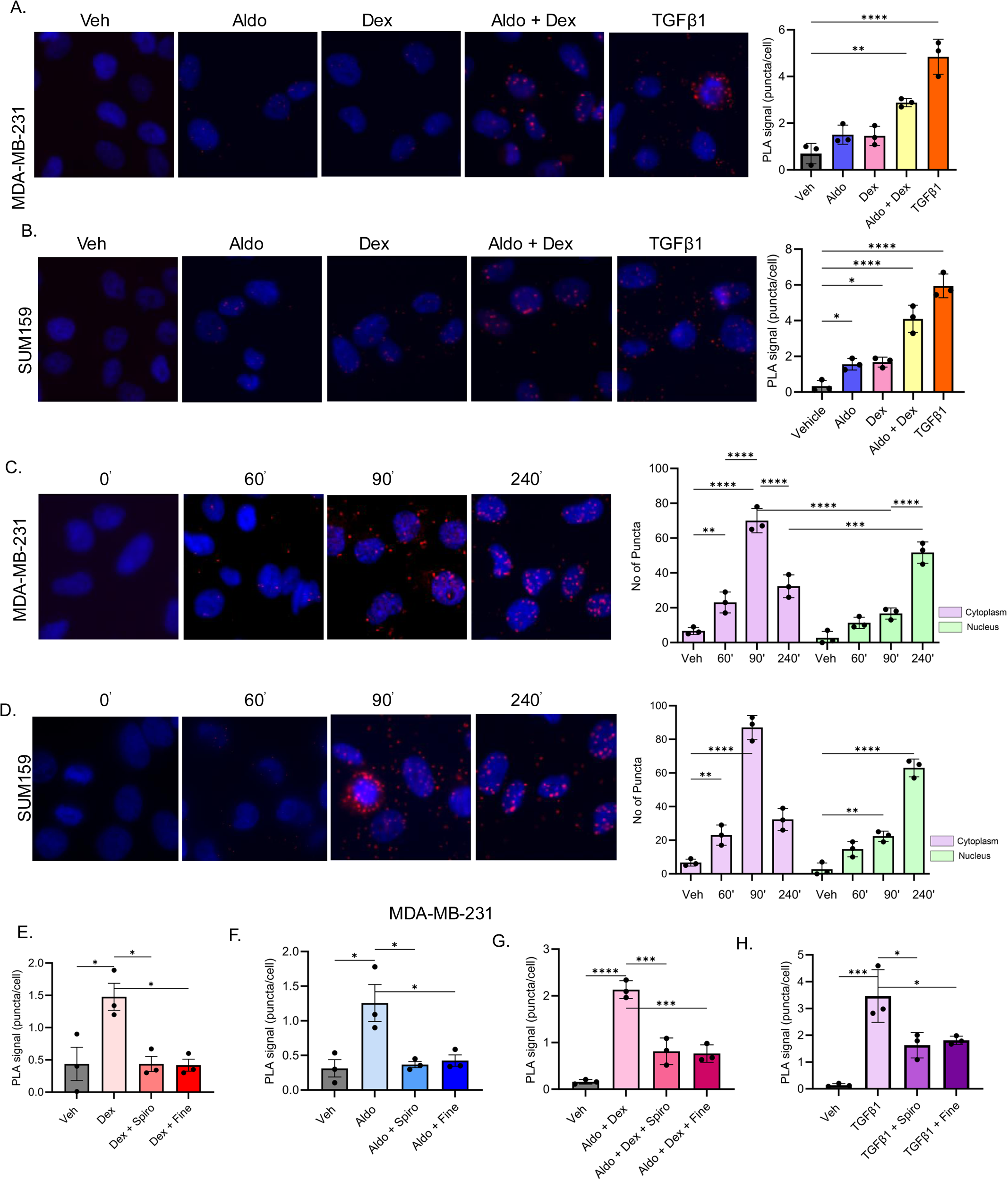
(A) Representative images of proximity ligation assays (PLA) showing an interaction between MR and GR in MDA-MB-231 cells treated with vehicle (veh),1 μM aldosterone (aldo), 1 μM dexamethasone (dex), 1 μM aldo plus 1 μM dex, or 10 ng/mL TGFβ1 for 90 min and quantification of the PLA signal (red puncta) using ImageJ. (B) Representative images of PLA showing an interaction between MR and GR in SUM159 cells treated as in panel A and quantification of the PLA signal (red puncta) using ImageJ. (C) Representative images of PLA showing an interaction between MR and GR in MDA-MB-231 cells with 10 ng/mL TGFβ1 for 0, 60, 90, and 240 min and quantification of the PLA signal (red puncta) using ImageJ. (D) Representative images of PLA showing an interaction between MR and GR in SUM159 cells treated with 10 ng/mL TGFβ1 for 0, 60, 90, and 240 min and quantification of the PLA signal (red puncta) in ImageJ. (E) Quantification of PLA signal (red puncta) in ImageJ from MDA-MB-231 cells treated with veh, 1 μM aldo for 90 min alone, or pretreatment with 10 μM spironolactone (spiro) or 10 μM finerenone (fine) for 60 min followed by 1 μM aldo treatment for 90 min. (F) Quantification of PLA signal (red puncta) in ImageJ from MDA-MB-231 cells treated with veh, 1 μM dex for 90 min alone, or pretreatment with 10 μM spiro or 10 μM fine for 60 min followed by 1 μM dex treatment for 90 min. (G) Quantification of PLA signal (red puncta) in ImageJ from MDA-MB-231 cells treated with veh, 1 μM aldo plus 1 μM dex for 90 min alone, or pretreatment with 10μM spiro or 10 μM fine for 60 min followed by 1 μM aldo plus 1 μM dex treatment for 90 min. (H) Quantification of PLA signal (red puncta) in ImageJ from MDA-MB-231 cells treated with veh, 10 ng/mL TGFβ1 for 90 min alone, or pretreatment with 10 μM spiro or 10 μM fine for 60 min followed by 10 ng/mL TGFβ1 for 90 min. The mean of three biological replicates is shown ± SD; *P < 0.05, **P < 0.01; ***P < 0.001; ****P < 0.0001.

Next, to determine if initial cytoplasmic TGFβ1-induced puncta ultimately become located to the nucleus, we performed a time course experiment. PLA was performed in MDA-MB-231 and SUM159 cells treated with 10 ng/mL TGFβ1 over a 4-hour time course. At a baseline of 0 hours and 60 min, minimal PLA signals were detected, indicating low levels of MR-GR interaction under resting conditions. Within 90 min of TGFβ1 treatment, a significant increase in PLA puncta was observed, and the majority of GR-MR-containing puncta were still localized to the cytoplasm. However, by 4 hours, a significant redistribution of PLA puncta to the nucleus was observed, and overall MR-GR interaction levels remained elevated compared to baseline (Figure 2C and 2D). These results indicate that TGFβ1 significantly induces the cytoplasmic formation of MR-GR complexes, and that prolonged exposure causes these complexes to translocate into the nucleus. The early cytoplasmic retention phase suggests a cytoplasmic role or preparatory interaction step, while the later nuclear accumulation supports a potential transcriptional function for the complex in response to TGFβ1 signaling.

To assess how MR antagonists modulate MR-GR complex formation dynamics in response to hormonal or TGFβ1 stimulation, we performed PLA after pre-treatment with either 10 μM spironolactone (sprio) or 10 μM finerenone (fine) antagonists. MDA-MB-231 cells were exposed to 1 μM aldo, 1 μM dex, the combination of both steroids, or to 10 ng/mL TGFβ1 for 90 min. Pre-treatment with either antagonist resulted in a marked reduction in MR-GR PLA signal relative to either hormone or TGFβ1 treatment alone (Figure 2E, 2F, 2G, 2H). These results align with their published pharmacological profile. Spironolactone, a steroidal MR blocker, is known to allow partial MR nuclear localization while impairing transcriptional activity through altered co-factor recruitment (54). Finerenone, a non-steroidal and more selective antagonist, more effectively impedes MR nuclear translocation and chromatin binding. Interestingly, both MR antagonists blocked the TGFβ1-induced cytoplasmic GR-MR interactions (Figure 2H). Collectively, these observations suggest that MR antagonists not only disrupt ligand-dependent MR signaling but also modulate receptor crosstalk and subcellular localization dynamics in response to both hormonal and cytokine (cellular stress) stimuli.

### TGFβ1-mediated MR and GR molecular complexes require activation of p38 MAPK and Ser134-GR phosphorylation

Since p38 MAPK is activated downstream of TGFβ1, we tested whether inhibiting p38 MAPK would impact MR-GR proximity. TNBC cells expressing wild-type GR and MR were pretreated with 10 μM of the selective p38 inhibitor SB203580 for 1 hour before 90 min of 10 ng/mL TGFβ1 stimulation. SB203580 alone did not affect baseline MR-GR PLA signal. Whereas upon TGFβ1 stimulation, we observed a pronounced increase in MR-GR complexes in the cytoplasm, in the presence of SB203580 + TGFβ1, the PLA signal significantly dropped compared to TGFβ1 alone (Figure 3A). Representative PLA images (Figure 3A) show markedly fewer red GR-MR-containing puncta in the perinuclear cytoplasm when p38 MAPK is inhibited. This finding establishes that p38 MAPK activity is required for efficient formation of MR–GR complexes in response to TGFβ1.

**Figure 3:**
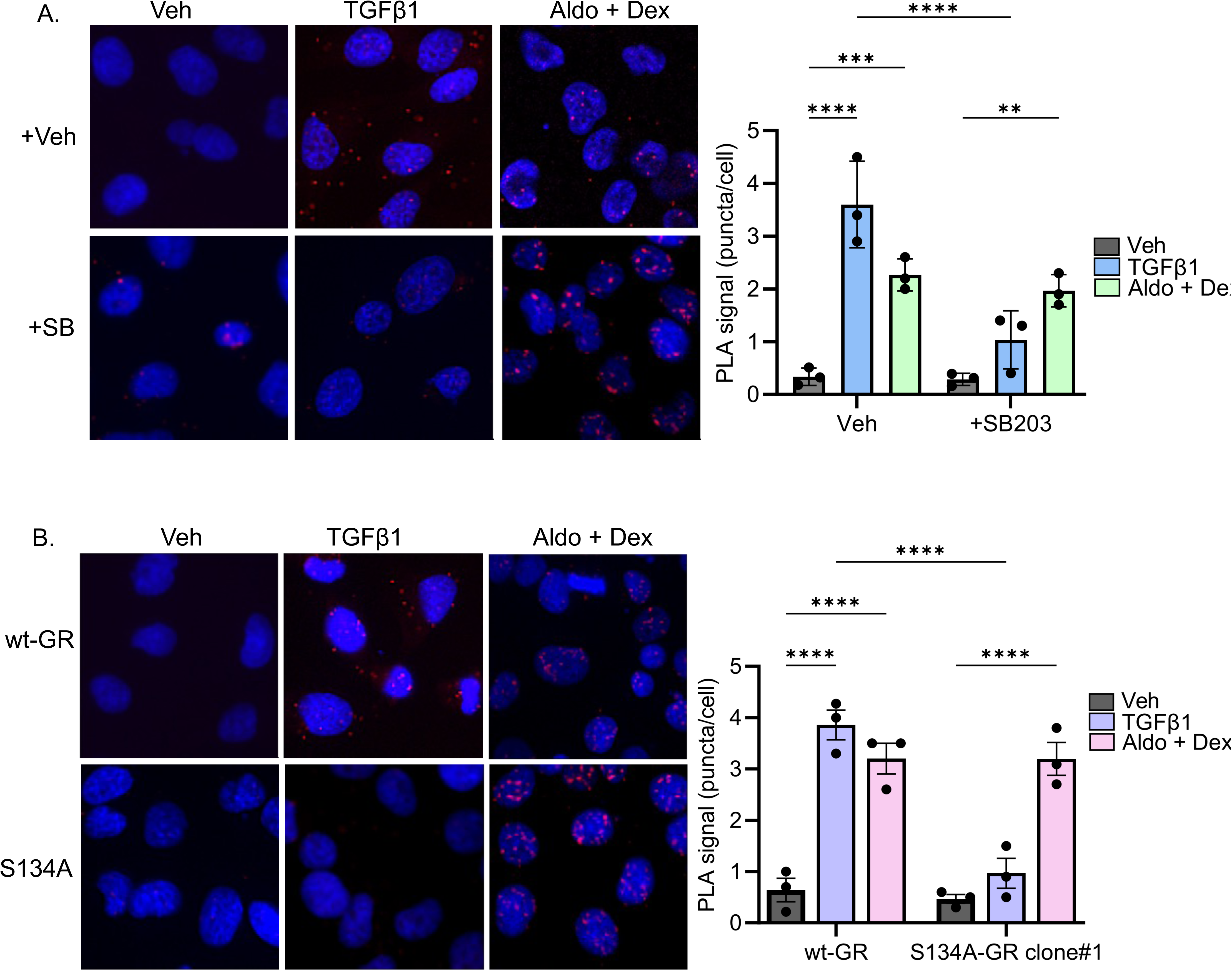
(A) Representative images of proximity ligation assays (PLA) showing interactions between MR and GR in MDA-MB-231 cells treated with vehicle (veh), 10 ng/mL TGFβ1 or 1 μM aldosterone (aldo) plus 1 μM dexamethasone (dex) for 90 min alone or pretreatment with 10 μM of SB203 for 30 min followed by treatment with veh, 10 ng/mL TGFβ1 or 1 μM aldo plus 1 μM dex for 90 min and quantification of PLA signal (red puncta) in ImageJ. (B) Representative images of PLA showing interactions between MR and GR in wt-GR and S134A-GR clone#1 MDA-MB-231 cells treated with veh, 10 ng/mL TGFβ1 or 1 μM aldo plus 1 μM dex for 90 min and quantification of PLA signal (red puncta) in ImageJ. The mean of three biological replicates is shown ± SD; *P < 0.05, **P < 0.01; ***P < 0.001; ****P < 0.0001.

Next, we asked if GR phosphorylation at Ser134 is necessary for MR–GR proximity, as Ser134 is a MAPK consensus site known to be regulated by p38. We first compared MR protein levels in the non-targeting control MDA-MB-231 sublines harboring wt-GR or in two independent Ser134 to alanine-GR CRIPSR knock-in clones (#1 and #2; reported in 16), confirming that all three lines have uniform MR and GR protein levels (Supplementary Figure 3A). As reported previously, the phosphorylation of GR by TGFβ1 at Ser134 is absent in the S134A-GR clones relative to cells expressing wt-GR (16). Treatment with 1 μM dex failed to induce pS134-GR, indicating that pSer134 is specifically a TGFβ1-mediated phosphorylation event (Supplementary Figure 3A).

Next, non-targeting control cells harboring either wt-GR or S134A-GR clone #1 and #2 were treated with 10 ng/mL TGFβ1 for 90 min and PLA was performed. While wt-GR cells exhibited cytoplasmic MR-GR puncta upon TGFβ1 stimulation, S134A-GR clone#1 cells exhibited a significantly lower PLA signal (Figure 3B). These data indicate that GR phosphorylation on Ser134 is a critical prerequisite for TGFβ1-induced MR–GR complex formation in the cytoplasm. Mechanistically, this suggests that p38 MAPK signaling and GR Ser134 phosphorylation are essential upstream steps for MR and GR to assemble under proinflammatory/profibrotic (TGFβ1) conditions typical of the TNBC tumor microenvironment. In contrast, MR-GR close-proximity nuclear interactions mediated by the combination of SR ligands aldo and dex was largely unaffected by mutation of GR Ser134 to alanine in clone#1 cells, which was verified in the S134A-GR clone#2 cell line (Supplementary Figure 3B). Together, these data indicate that, unlike TGFβ1–driven MR-GR complex formation, the classical ligand-induced MR–GR association requires neither p38 MAPK signaling nor GR Ser134 phosphorylation. In sum, aldo and dex can drive MR and GR into close-proximity/complexes independently of the kinase-dependent/ligand-independent pathway used by TGFβ1 signaling.

### Pharmacological inhibition of MR attenuates aldosterone and TGFβ1-induced migration and tumorsphere formation

Previously, we reported that cytokines from the tumor microenvironment, including TGFβ1, promote GR-mediated cell migration in scratch wound and transwell assays (16). Additionally, we showed that dexamethasone (dex) promoted biphasic cell migration in TNBC models, such that a 15 min acute pre-treatment with dex inhibited cell migration in scratch assays, while a treatment for 6 hours stimulated robust cell migration (16). Aldosterone (aldo), the physiological ligand for MR, has been shown to promote the migration of myoblasts and vascular smooth muscle cells (55, 56); however, the effects of MR agonists on the migration of TNBC cells have not been explored. For our study, MDA-MB-231 and SUM159 cells were plated in transwell chambers and treated with 1 μM aldo, 1 μM dex or 10 ng/mL TGFβ1. TNBC cells were allowed to migrate for 6 hours using 10% FBS as the chemoattractant. Dex and aldo stimulated the migratory capacity of both MDA-MB-231 (Figure 4A and 4B) and SUM159 cells (Figure 4D and 4E). In a previous study we showed that the TGFβ1 pathway was among the most significantly upregulated pathways upon chronic exposure of MDA-MB-231 cells to either dex or TGFβ1 treatment (16). As expected, we observed an increase in cell migration upon TGFβ1 treatment in transwell migration (Figure 4C, 4F). Notably, TNBC cells treated with the MR agonist aldo behaved similarly to cells treated with either dex or TGFβ1, exhibiting ∼2.5-3 fold increased cell migration over the same time frame (Figure 4A, and 4E).

**Figure 4:**
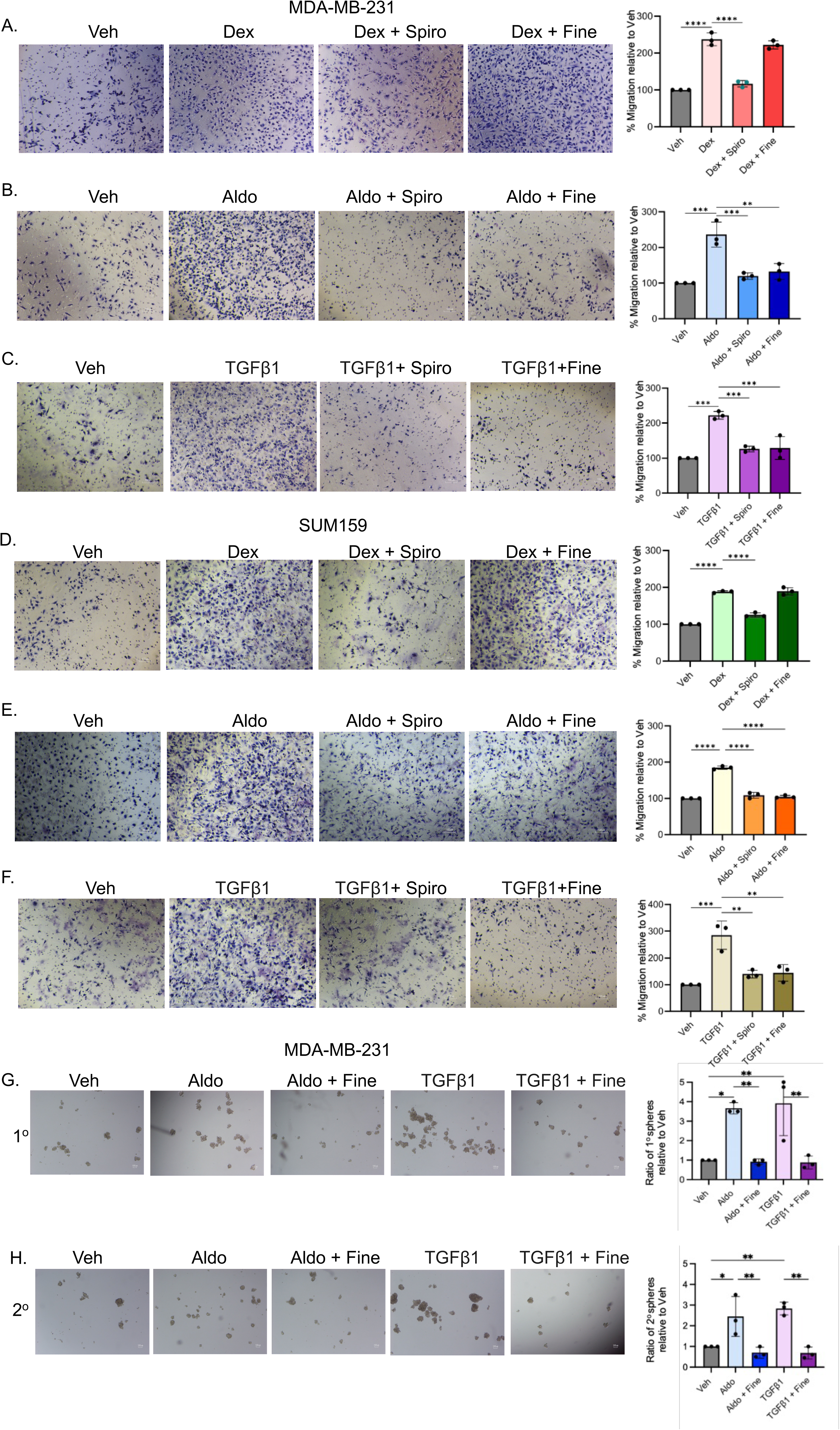
(A-F) Representative images of transwell assays and the percentage of migrated cells, relative to vehicle (veh) controls (set to 100%). (A) MDA-MB-231 cells were treated with 1 µM dexamethasone (dex), 1 µM dex plus 10 µM spironolactone (spiro), or 1 µM dex plus 10 µM finerenone (fine). (B) MDA-MB-231 cells were treated with 1 µM aldosterone (aldo), 1 µM aldo plus 10 µM spiro, and 1 µM aldo plus 10 µM fine. (C) MDA-MB-231 cells were treated with 10 ng/mL TGFβ1, 10 ng/mL TGFβ1 plus 10 µM spiro, and 10 ng/mL TGFβ1 plus 10 µM fine. (D) SUM159 cells were treated with 1 µM dex, 1 µM dex plus 10 µM spiro, or 1 µM dex plus 10 µM fine. (E) SUM159 cells were treated with 1 µM aldo, 1 µM aldo plus 10 µM spiro, and 1 µM aldo plus 10 µM fine. (F) SUM159 cells were treated with migrated 10 ng/mL TGFβ1, 10 ng/mL TGFβ1 plus 10 µM spiro, or 10 ng/mL TGFβ1 plus 10 µM fine. (G-H) Representative images of (G) primary or (H) secondary tumorspheres generated from MDA-MB-231 cells treated with veh, 1 μM aldo, 10 ng/ml TGFβ1, 1µM aldo plus 10 µM fine or 10 ng/mL TGFβ1 plus 10 µM fine; bar graphs show the ratio of sphere counts relative to the vehicle control (set to 100%). The mean of three biological replicates is shown ± SD; *P < 0.05, **P < 0.01; ***P < 0.001; ****P < 0.0001.

We next examined the requirement of MR activity in dex, aldo and TGFβ1-induced TNBC cell migration by pre-treating cells with two MR antagonists, spironolactone (spiro) and finerenone (fine) for one hour. Following pretreatment, cells were exposed to aldo, dex, or TGFβ1, and allowed to migrate for 6 hours using 10% FBS as the chemoattractant in the lower chamber. Spiro, a first-generation steroidal MR antagonist, at 10 μM effectively inhibited TNBC cell migration induced by aldo, dex or TGFβ1 (Figure 4A, 4B, 4C, 4D, 4E and 4F). In contrast, 10 μM of fine, a third-generation non-steroidal and highly selective MR antagonist, preferentially blocked migration in response to either aldo or TGFβ1, but not to dex (Figure 4A, 4B, 4C, 4D, 4E and 4F). These differential effects may be attributed to the off-target GR antagonism by spiro, which, although primarily targets MR (Ki ≈ 24 nM), also exhibits weak glucocorticoid receptor (GR) binding (Ki ≈ 620 nM) (54). In contrast, fine, which has a much lower affinity for GR (Ki > 10,000 nM) than spiro, failed to interfere with dex signaling, consistent with its high MR selectivity. Collectively, our data suggest that cooperation between GR and MR mediates TGFβ1-induced cell migration, while in response to their cognate steroid hormone agonists, both GR and MR can independently stimulate increased TNBC cell migration.

To test the requirement of MR in TNBC cancer stem-like phenotypes, tumorsphere assays were used in which TNBC cells were plated at limiting dilution in ultra-low attachment dishes in the presence of methylcellulose to prevent aggregation (57). Interestingly, 10 µM fine inhibited both primary and secondary tumorsphere formation mediated by aldo or TGFβ1 (Figure 4G, and 4H). These results demonstrate that pharmacological inhibition of MR with finerenone significantly reduces both the initiation (primary spheres) and self-renewal of tumorspheres (secondary spheres) in TNBC cell lines. The suppression of secondary sphere formation indicates that MR signaling is critical not only for tumor-initiating cell survival, but also for their capacity to regenerate spheres from single cells under stem cell-selective conditions.

### MR knockdown transcriptionally downregulates both MR and p-GR target genes

To verify that loss of MR function impacts transcriptional activity in TNBC cells, we created stable shRNA-mediated MR knockdown in both MDA-MB-231 and SUM159 lines, as well as non-targeting vector controls (shGFP). MR knockdown was confirmed by Western blotting and qPCR (Figure 5A, 5B and 5C). Densitometry values are shown in Supplementary Figure 4A and 4B. To study the impact of reduced MR expression on transcriptional regulation of known MR and/or GR target genes, we compared mRNA levels of canonical and non-canonical target genes. Canonical MR target genes are typically expressed in aldosterone (aldo)-sensitive epithelial tissues, such as the kidney, colon, and salivary glands, and are crucial for maintaining sodium and water balance. Key MR canonical targets include SGK1 (serum and glucocorticoid-regulated kinase 1) and several epithelial sodium channel (ENaC) subunits (SCNN1A, SCNN1B, SCNN1G) (58, 59). These genes mediate the classical physiological effects of aldo, particularly in blood pressure regulation and electrolyte homeostasis.

**Figure 5:**
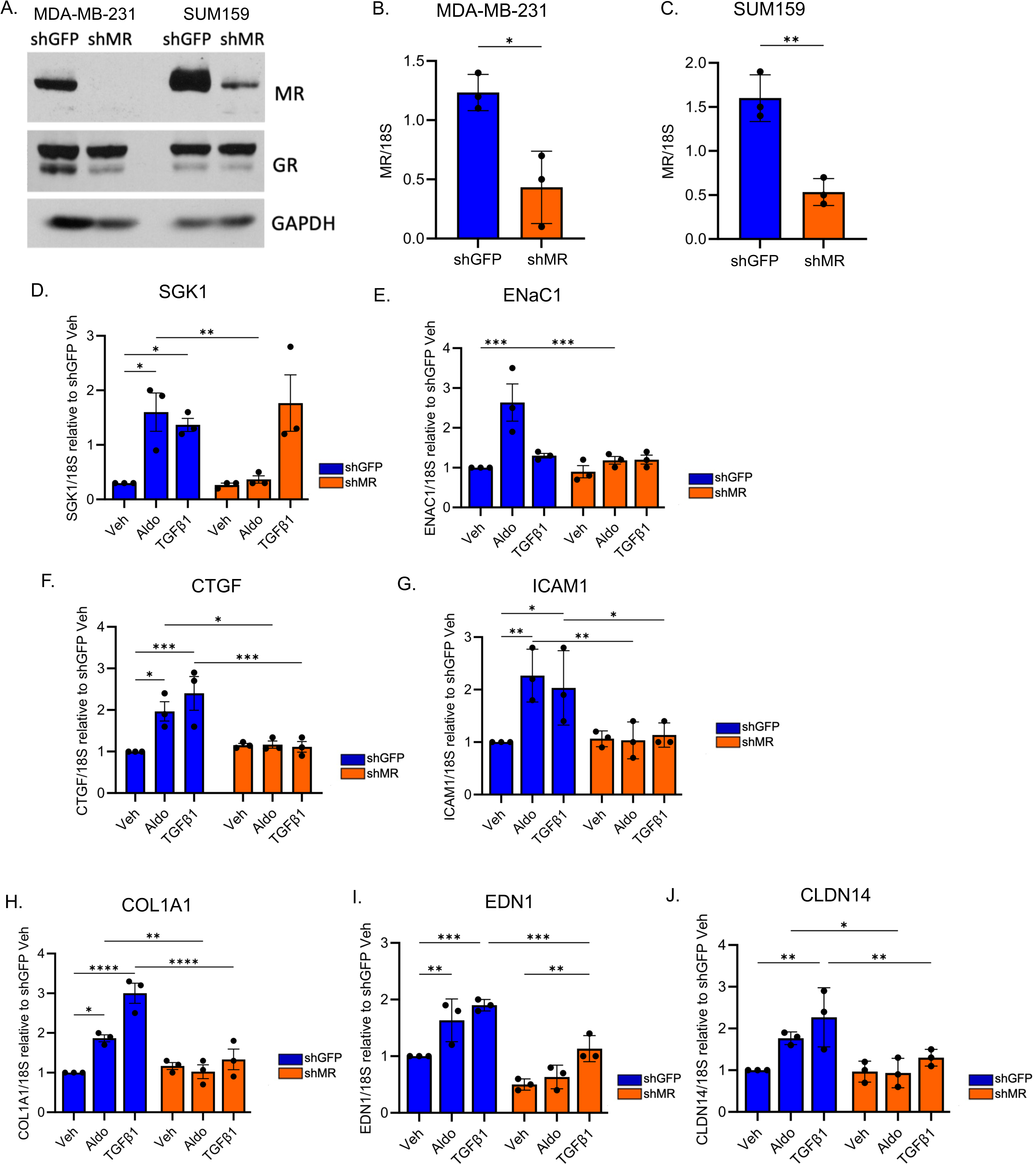
(A) Western blot of MR and GR protein levels in MDA-MB-231 and SUM159 cells following transduction with shGFP (negative control) or shMR lentivirus particles. (B-C) MR mRNA levels in (B) MDA-MB-231 and (C) SUM159 shGFP and shMR cells were assessed by qRT-PCR following normalization to 18S mRNA. (D-J) mRNA levels of MR canonical and non-canonical genes, including (D) SGK1, (E) ENaC1, (F) CTGF, (G) ICAM1, (H) COL1A1, (I) EDN1 and (J) CLDN14, were assessed by RT-qPCR in shGFP (blue bars) and shMR MDA-MB-231 (orange bars) cells treated with vehicle (veh), 1 µM aldosterone (aldo) or 10 ng/mL TGFβ1 for 6 hours. The mean of three biological replicates is shown ± SD. (*, P < 0.05, **, P < 0.01, ***, P < 0.001, ****, P < 0.0001).

Aldo induced upregulation of SGK1 and SCNN1A in shGFP MDA-MB-231 cells, but not in shMR cells (Figure 5D and 5E). In contrast, TGFβ1 treatment did not alter the expression of these genes, supporting the specificity of MR signaling in their regulation (Figure 5D and 5E). Likewise, whereas shGFP SUM159 cells also upregulate SGK1 and ENaC1 when stimulated by aldo and TGFβ1, induction is suppressed in shMR SUM159 cells (Supplementary Figure 4C and 4D). These findings reinforce the concept that ENaC1 and SGK1 are classical, directly-regulated targets of MR, but are not responsive to TGFβ1–mediated signaling, which is more typically associated with non-canonical MR functions such as fibrosis or inflammation.

Non-canonical MR target genes are those regulated by the receptor through indirect or context-dependent mechanisms, often outside classical epithelial tissues. Unlike canonical targets like ENaC and SGK1, the non-canonical genes do not typically involve direct MR binding to hormone response elements (29). Instead, they are influenced by MR activity through crosstalk with signaling pathways such as TGF-β, NF-κB, or MAPK, and are often associated with inflammation, fibrosis, oxidative stress, and tissue remodeling (29). We observed a significant upregulation in the levels of connective tissue growth factor (CTGF), Intercellular Adhesion Molecule-1 (ICAM1), and alpha 1 chain of type I collagen (COL1A1) (Figure 5F, 5G and 5H). Both aldo and TGFβ1 treatment led to upregulation of these non-canonical MR target genes, which was abolished upon MR knockdown. Similarly, aldo-induced upregulation of CTGF in shGFP SUM159 cells was absent in shMR SUM159 cells (Supplementary Figure 4E). Aldo and TGFβ1-induced upregulation of ICAM1 in shGFP SUM159 cells was absent in shMR SUM159 cells (Supplementary Figure 4F). These findings highlight a broader, non-canonical role for MR in modulating gene expression beyond classical aldo-responsive targets. This may reflect a non-classical role for MR in facilitating transcriptional responses to TGFβ1, possibly through receptor crosstalk, co-regulator recruitment, and/or chromatin remodeling.

Notably, we have shown that COL1A1 is a known TGFβ1-induced p-GR target gene, and that when the GR Ser134 is mutated to alanine, TGFβ1-induced upregulation of COL1A1 was lost (16). Therefore, we compared the transcriptional regulation of additional p-GR target genes by MR after knockdown. Interestingly, we observed similar aldo and TGFβ1-induced upregulation of EDN1 and CLDN14, which is lost upon MR knockdown (Figure 5I and 5J). Extending these findings, MR-dependence of COL1A1, EDN1 and CLDN14 expression highlights a cooperative role for MR in supporting p-GR-driven transcription and a functional interplay between MR and p-GR signaling pathways. Collectively, these results support a model in which MR and p-GR cooperate to regulate a subset of non-canonical target genes in response to proinflammatory (cytokine), profibrotic or hormonal stimuli in TNBC, thereby expanding the known set of MR’s regulatory functions beyond classical mineralocorticoid-responsive pathways.

### MR mediates aldosterone- as well as TGFβ1-induced TNBC cell migration and tumorsphere formation via a p38 MAPK-dependent mechanism

Next, to directly test the requirement of MR in aldosterone (aldo) and TGFβ1-induced TNBC cell migration we subjected MDA-MB-231 and SUM159 shMR cells as well as their paired shGFP controls to transwell migration assays. TNBC cells were treated with 1 µM aldo or 10 ng/mL TGFβ1 and allowed to migrate for 6 hours with 10% FBS as the chemoattractant in the lower chamber. The results showed that aldo and TGFβ1 induced robust cell migration in shGFP, but not in shMR MDA-MB-231 cells (Figure 6A). SUM159 cells showed a similar phenotype wherein aldo and TGFβ1 induced migration in shGFP cells, which was suppressed in cells expressing shMR (Figure 6B). To rule out off-target shRNA MR-mediated effects and to minimize compensation due to either antibiotic selection or the stable expression of the MR shRNA, MDA-MB-231 cells were also subjected to transient siRNA-mediated MR knockdown (Supplementary Figure 5A and 5B). When stimulated with TGFβ1 or aldo, the migratory capacity of TNBC cells transfected with MR siRNA was significantly hindered compared to the cells transfected with scramble (scr) siRNA (Supplementary Figure 5C).

**Figure 6:**
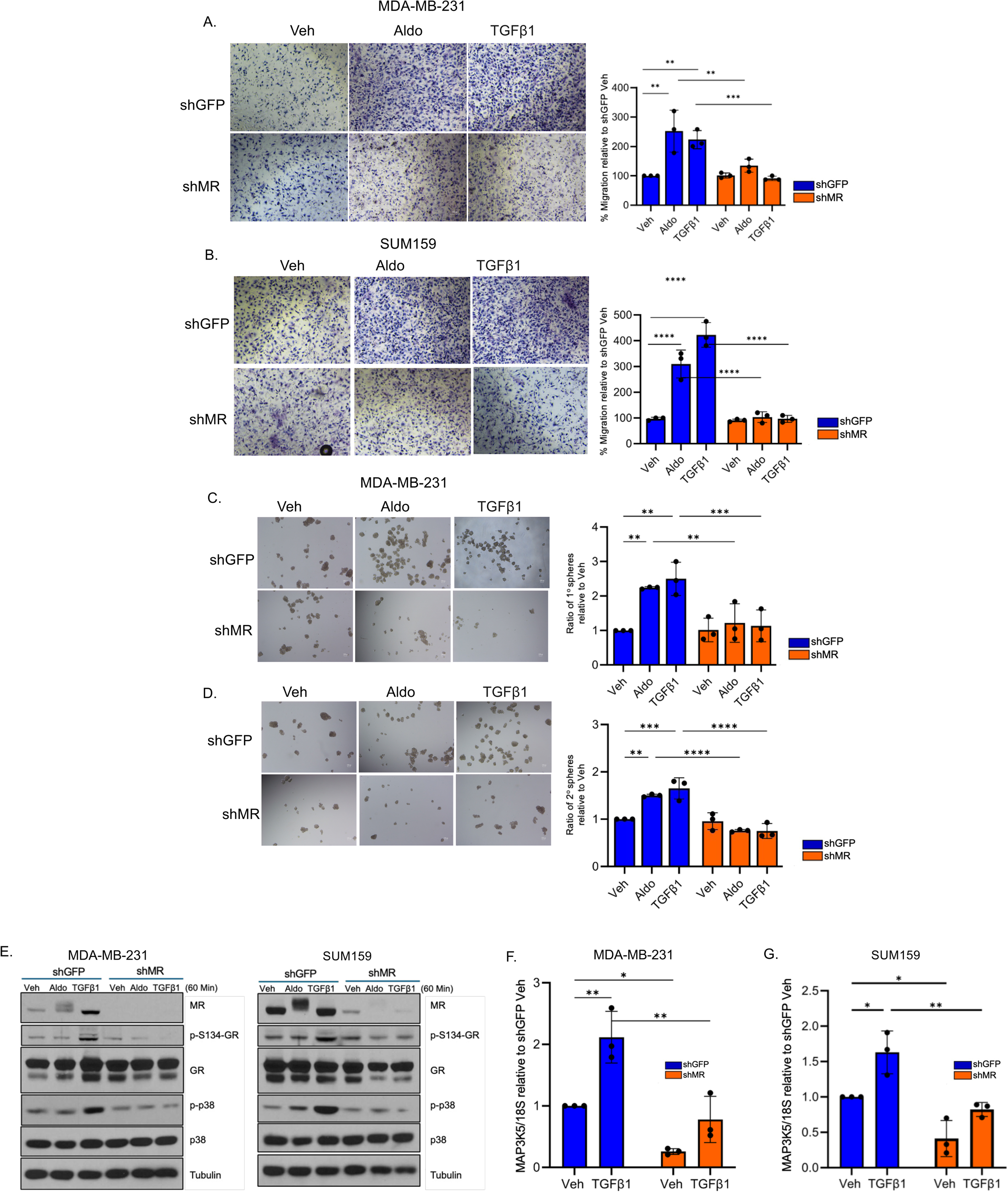
(A-B) Representative images of transwell migration assays and the percentage of migrated cells in shGFP and shMR MDA-MB-231 cells relative to vehicle (veh) treatment (set to 100%). (A) MDA-MB-231 or SUM159 (B) cells were treated with 1 μM aldosterone (aldo) or 10 ng/mL TGFβ1 and allowed to migrate towards 10% FBS for 6 hours. (C-D) Representative images of (C) primary or (D) secondary tumorspheres generated from shGFP and shMR MDA-MB-231 cells, expressed as a ratio relative to veh, treated cells (set to 1.0), following treatment with 1 μM aldo or 10 ng/ml TGFβ1. The Mean of three biological replicates is shown ± SD: *, P < 0.05, **, P < 0.01, ***, P < 0.001, ****, P < 0.0001. (E) Representative western blot analysis of MR, pS134-GR, total GR, p-p38, total p38, and tubulin (loading control) levels in MDA-MB-231 and SUM159 shGFP and shMR cells treated with 1 μM aldo or 10 ng/ml TGFβ1 for 60 minutes. (F-G) Bar graphs of the mRNA levels of MAP3K5, normalized to 18S mRNA levels of vehicle-treated cells, as compared by RT-qPCR in (F) shGFP and shMR MDA-MB-231 or (G) SUM159 cells treated with either veh or 10 ng/mL TGFβ1 for 24 hours. The Mean of three biological replicates is shown ± SD: *, P < 0.05, **, P < 0.01, ***, P < 0.001, ****, P < 0.0001.

Further, to test the requirement of MR for cancer stem cell biology, tumorsphere assays were performed in which shGFP and shMR MDA-MB-231 cells were plated at limiting dilution in ultra-low attachment dishes. The results showed that in the presence of 1 µM aldo or 10 ng/mL TGFβ1, shMR cells formed significantly fewer primary and secondary tumorspheres relative to shGFP controls (Figure 6C and 6D). These results were confirmed as measured by sphere counts. (Figure 6C and 6D).

To assess the mechanism of reduced TGFβ1-induced migration and tumorsphere formation in MR KD cells at the level of cellular stress pathway signaling, we probed for phosphorylation of GR at Ser134 and p38 MAPK in shGFP and shMR cells. Indeed, TGFβ1-induced (i.e. p38-dependent) phosphorylation of GR Ser134 appears to be disabled in cells expressing shMR relative to cells expressing shGFP (Figure 6E). Densitometric analyses of aggregated Western blot data from numerous independent biological replicates is shown (Supplementary Figure 5D and 5E). Similarly, MR knockdown resulted in greatly decreased TGFβ1-induced p38 MAPK activation (as measured by phosphorylation of kinase active site residues) with no change in total p38 MAPK expression levels (Figure 6E). We previously reported that TNBC cells expressing S134A-GR exhibited reduced levels of cytokine-activated p38 MAPK with no effect on total p38 levels (16). Further, cells expressing S134A-GR exhibited downregulation of basal MAP3K5 mRNA and protein expression. MAP3K5 (ASK1) is a protein kinase with cancer relevance that interacts with 4-3-3ζ to facilitate activation of downstream kinases MEK3/6 and p38 MAPK at both the mRNA and protein levels (16). As our data demonstrate that MR KD phenocopies S134A-GR, we assessed mRNA levels of MAP3K5 in shGFP and shMR MDA-MB-231 and SUM159 cells treated with 10 ng/mL TGFβ1 for 6 hours or 24 hours. We confirmed that MR KD cells exhibit a downregulation of basal MAP3K5 levels relative to the control cells. While MAP3K5 mRNA was not appreciably induced by TGFβ1 at 6 hours (Supplementary Figure 5F and 5G), we observed upregulation of MAP3K5 mRNA at 24 hours of TGFβ1 treatment that was significantly hindered in MR KD cells (Figure 6F and 6G). These data suggest that MR is an important regulator of MAP3K5 expression, a limiting kinase upstream of p38 MAPK that is required for activation of the p38 MAPK module and subsequent GR Ser134 phosphorylation in response to cellular stress stimuli.

### MR signaling contributes to TNBC metastasis to lungs *in vivo*

To test the impact of loss of MR signaling to metastasis *in vivo*, shGFP and shMR MDA-MB-231 cells were injected into the tail vein of female NSG mice. Lungs were harvested 21 days later, paraffin-embedded and sectioned, and then stained with H&E. QuPath software was used to mark all areas of tumor metastasis, and the area occupied by metastases/total lung area was calculated as a percentage. The mean area occupied by metastases in shGFP mice was 44.2%, which was reduced ∼2-fold in shMR-injected mice to 22.7% (Figure 7A and 7B).

**Figure 7:**
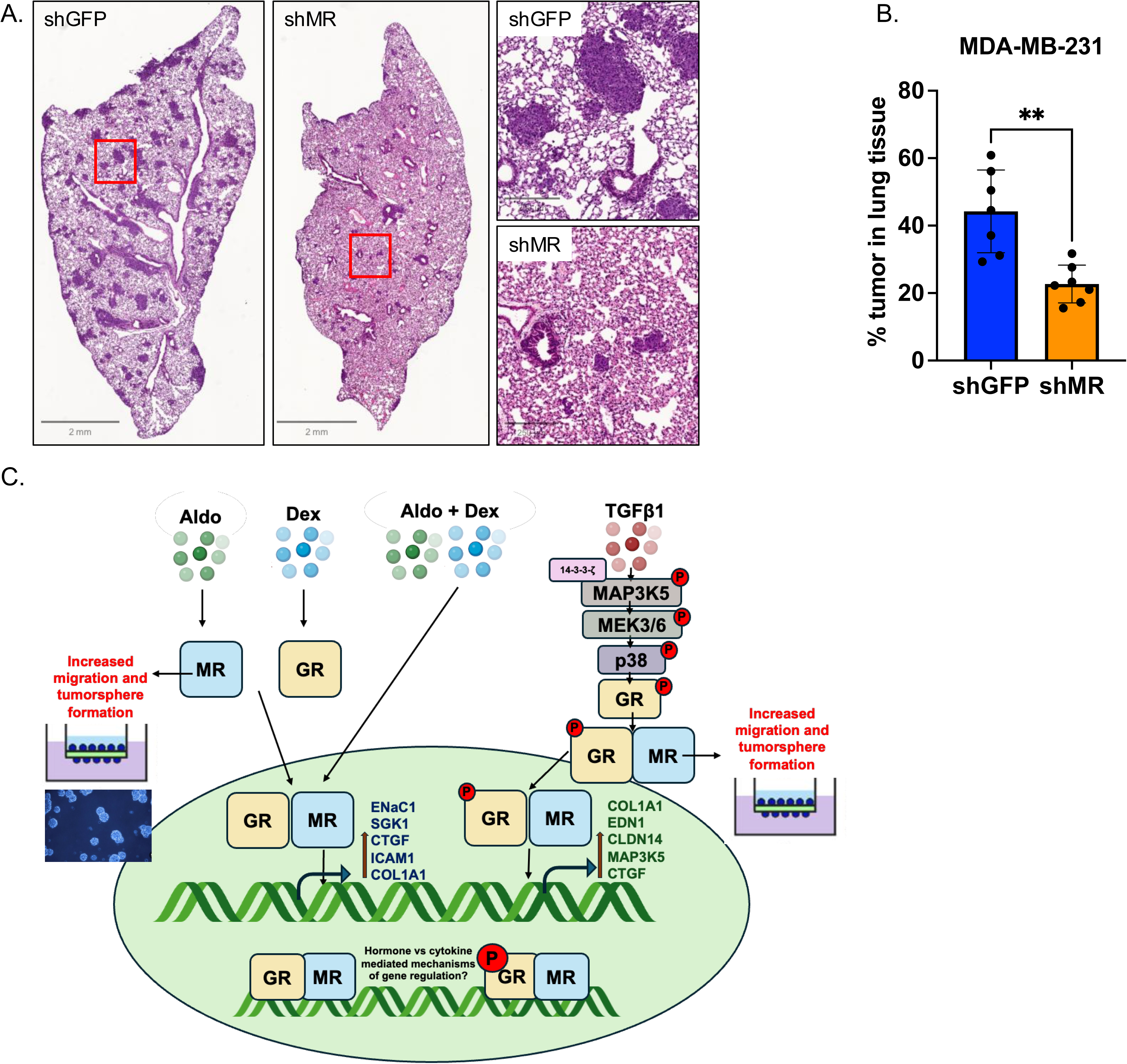
(A) Representative H&E-stained lung tissues following tail-vein injection of shGFP and shMR MDA-MB-231 cells. (B) Percentage (%) tumor area in lung calculated from H&E sections. ** P < 0.01. (C) Summary of Findings - Aldosterone (aldo), dexamethasone (dex) or a combination of both promotes MR and GR proximity in the nucleus. MR activation by aldo then upregulates genes such as ENaC1, SGK1, CTGF, ICAM1 and COL1A1. On the other hand, TGFβ1 promotes MR and GR proximity association in the cytoplasm, mediated by phosphorylation of p38 MAP kinase and GR at Ser134 which translocates to the nucleus. MR activation by TGFβ1 upregulates fibrotic genes such as CTGF, ICAM1 and COL1A1 and p-Ser134-GR target genes such as EDN1, CLDN14 COL1A1 and MAP3K5. Future studies are focused on identifying differences between hormone and TGFβ1-mediated mechanisms of gene regulation by MR.

## Discussion

The role of MR signaling in TNBC remains largely unexplored. In this study, we examined its potential involvement in cancer hallmarks/metastatic traits associated with TNBC. Collectively, our data provide the first *in vitro* and *in vivo* evidence that targeting MR/p-GR-may be a useful strategy to block TNBC cell migratory, stemness, and metastatic phenotypes that are mediated by stress-responsive steroid hormones such as aldosterone or cortisol, as well stress-induced cytokines, including TGFβ1 produced by the tumor microenvironment, especially in weakly or non-proliferative circulating tumor cell sub-populations that are relatively insensitive to classical cytotoxic chemotherapies. Our study defines a novel and actionable signaling axis in TNBC in which MR forms a critical signaling co-complex with phosphorylated GR to promote pro-migratory, cancer stem-like, and metastatic phenotypes in response to TGFβ1 (Figure 7C). Moreover, these aggressive phenotypes mediated by the MR/p-GR co-complex can be effectively inhibited by FDA-approved MR antagonists. These MR antagonists, particularly finerenone, may provide a more selective and durable up-front treatment option for patients with TNBC since systemic chemotherapy, the primary standard-of-care for TNBC, often fails to prevent rapid development of chemoresistance and early relapse (60).

Targeting steroid hormone receptors other than ER and PR is gaining increased attention. There are distinct subtypes of TNBC that express either androgen receptor (AR) or GR (61, 62). GR, PR, and MR are closely related hormone receptors and recognize similar DNA binding sites in chromatin (63). We have previously shown that p38-dependent phosphorylation of GR at Ser134 correlates with poor prognosis in TNBC (16). We now show that MR levels also correlate with poorer survival of TNBC patients, and that MR/pSer134-GR complexes form in response to either co-treatment with aldo and dexamethasone (dex) or exposure to TGFβ1. MR activity was necessary to promote cell migration, primary and secondary tumorsphere formation, and to induce expression of canonical and non-canonical MR gene expression programs in TNBC cells. We also determined that knockdown of MR reduced the phosphorylation of GR at Ser134 and p-p38, explaining why MR antagonists or MR loss of function phenocopies the mutation of GR Ser134 to alanine in TNBC cells, which we previously reported (16). Like TNBC models harboring phospho-mutant S134A GR, TNBC cells failed to colonize mouse lungs *in vivo* following MR knockdown. These results demonstrate that the MR-pGR complex is a clinically relevant prognostic and functional driver of TNBC.

Beyond its classical epithelial roles in kidney and colon cells, MR has been implicated in pathological tissue remodeling and fibrosis in cardiovascular disease models (64). In murine models of chronic blood pressure overload and myocardial infarction, deletion or pharmacologic inhibition of MR (NR3C2) attenuated left ventricular dilation, cardiac hypertrophy, and heart failure progression, whereas MR overexpression in cardiomyocytes induced fibrosis, adverse remodeling, and pro-arrhythmogenic effects (65). These findings collectively illustrate that MR activation can drive pathological remodeling processes beyond fluid homeostasis, consistent with our findings that MR also mediates pro-migratory and stem-like programs in TNBC through its interaction with GR and p-GR.

In a breast cancer study exploring the expression of MR and GR, along with expression of 11β-hydroxysteroid dehydrogenase enzymes (11βHSD1/2), across cancer progression (66), MR expression levels, particularly nuclear MR, increased from atypical ductal hyperplasia to invasive ductal carcinoma (IDC-NST). GR expression also increased (66). Notably, the 11βHSD1/2 ratio is linked to modulation of local cortisol availability and was elevated in ductal carcinoma in situ (DCIS), suggesting glucocorticoids may predominantly activate MR and/or MR-GR in early breast cancer. MR expression showed subtype-specific correlations with proliferation markers and potential crosstalk with PR (i.e. in ER+ tumors), implying a complex hormonal regulatory role that may differ across breast cancer subtypes (67).

We have examined MR’s potential involvement in cancer hallmarks/metastatic traits associated with TNBC. First, analysis of public datasets revealed that MR mRNA levels are enriched in TNBC (ER-/HER2-) compared to other subtypes. Notably, higher MR expression was also linked to poorer overall survival in these patients. These data mining findings suggest that MR is more highly expressed in TNBC and may contribute to its aggressive behavior, providing a rationale for investigating its functional significance in TNBC progression to metastasis. It has been demonstrated previously that GR phosphorylation at the N-terminal Ser134 p38 MAPK site is crucial for TGFβ1 induced-TNBC cell migration, invasion, anchorage-independent growth, tumorsphere formation, and metastasis in mouse models (16). RNA-sequencing data from these studies also showed that p-S134-GR is required for expression of genes that promote MAP kinase activity in TNBC and MAPK signaling has been heavily implicated in EMT, stemness and chemoresistance, which are largely non-proliferative mechanisms that underlie metastatic breast cancer progression (68). In the present study, we used two distinct and robust *in vitro* techniques as well as *in vivo* (tail vein) studies to define the importance of MR in advanced cancer phenotypes as summarized in Figure 7C, including the transwell migration assay and the tumorsphere assay. In transwell assays, we observed an increase in migratory capacity of two TNBC cell lines towards the chemoattractant (10% FBS) in the presence of aldo or TGFβ1. The observed inhibition of aldo- and TGFβ1-induced migration with pretreatment of finerenone (fine), a fourth generation MR antagonist, suggests MR as a critical mediator of cytokine-driven TNBC cell migration. Fine is a non-steroidal, selective MR antagonist with high affinity and specificity for MR over GR. Unlike classical steroidal MR antagonists such as spironolactone (spiro), fine does not exhibit significant off-target activity at androgen receptor, progesterone receptor, or GR, thereby minimizing side effects related to hormonal cross-reactivity. Importantly, in our transwell assays, fine does not block the effects of dex, a potent GR agonist, further confirming its receptor selectivity and supporting the conclusion that the actions of Dex (i.e., at low doses) are mediated primarily through GR rather than MR. This pharmacologic specificity makes fine a valuable tool for dissecting MR-specific signaling pathways without the confounding effects on GR-mediated responses. Additionally, MR has nine amino acid residues (Ser or Thr) in its primary structure that are strong consensus sites for proline-directed kinases such as MAPKs; we predict that this allows MR to sense relatively small changes in local stress signals from cytokines such as TGFβ1 or other factors rich within the tumor microenvironment (69). Analysis of these potential phosphorylation sites within the MR N-terminus and their interplay with GR and p-GR is a topic for future study.

We also show that fine blocks both primary and secondary tumorsphere formation that is induced by aldo or TGFβ1. All treatments were compared in both primary and secondary tumorspheres to provide a functional framework to assess the impact of MR signaling on cancer stem-like cell expansion or potential. The primary tumorsphere assays informed us about initial sphere formation potential whereas the secondary tumorsphere assay reveals the sustainability of the self-renewing population that expanded in response to MR activity. These results underscore the importance of MR in mediating the amplification and maintenance of cancer stem-cell like populations in TNBC models. Accordingly, we observed reduced TNBC lung metastases *in vivo* following MR knockdown relative to parental controls using the mouse tail vein injection assay, which model late events in the metastasis cascade (i.e., TBNC cell survival in the circulation, extravasation from the vasculature, and lung colonization/tumor formation).

In RT-qPCR studies, aldo-induced upregulation of genes such as ENaC1 and SGK1 was significantly abolished in MR-KD cells confirming an MR-mediated activation of ENaC1 and SGK1. MR plays a central role in the pathogenesis of fibrosis across multiple organs, including the heart, kidney, vasculature, and lungs (67). Beyond its classical role in sodium homeostasis, MR activation in non-epithelial cells such as fibroblasts, macrophages, and endothelial cells leads to pro-inflammatory and pro-fibrotic gene expression, independent of blood pressure regulation. We observed a significant loss in aldo- and TGFβ1-mediated upregulation of CTGF, ICAM1 and COL1A1 genes in MR-KD cells. It has been reported that MR activation enhances the expression of CTGF, a connective tissue growth factor that leads to fibroblast activation and collagen deposition (70–72). Aldo–MR signaling also promotes COL1A1 transcription in cardiac fibroblasts, renal mesangial cells, and vascular smooth muscle cells. COL1A1 encodes the alpha-1 chain of type I collagen and is a core extracellular matrix component in fibrotic tissues (55, 73). In TNBC, high expression of COL1A1 is a poor prognostic indicator that correlates with increased lymph node metastasis, larger tumor size, and shortened patient survival (74–77). ICAM1, an adhesion molecule that has been shown to be expressed in endothelial and smooth muscle cells, is regulated in an MR-dependent manner and contributes to vascular inflammation and immune infiltration, processes integral to the pathogenesis of fibrosis (78, 79). In TNBC, elevated expression of ICAM1 is linked to promoting bone and lung metastasis though TGF-β/SMAD pathways and also contributes to therapy resistance (80, 81). Interestingly, our previous RNA sequencing data in S134A-GR+ TNBC cells, showed that COL1A1 is a TGFβ1-induced pS134GR target gene. We observed a loss in regulation of EDN1 and CLN14 in MR-KD models, also previously identified TGFβ1-induced pS134-GR target genes. The loss of regulation of p-GR target genes in MR-deficient cells points to a previously unrecognized required role for MR in integrating TGF-β (cellular stress) and pS134-GR signaling (host or life stress), with potential implications in tissue remodeling, fibrosis, and ultra-sensitive or decisive responses to cellular stress stimuli in steroid hormone-responsive pathologies including TNBC or other cancers characterized by a highly pro-inflammatory TME.

TGFβ1 expression has been observed in 52.5% of TNBC tissue samples relative to non-TNBC cases (27.5%) (82). Additionally, patients with a higher TGFβ1 expression have a poorer 5-year disease-free survival rate (83). Further, in cell line models, TGFβ1 and β2 expression levels are higher in TNBC cells compared to non-TNBC cells, and the TNBC cells also highly express EMT-related gene signatures (83). Previous studies in our lab concluded that pS134-GR is a critical effector of migratory and stem-cell like behaviors linked to TGFβ1 signaling in TNBC models (16). The biological phenotype of decreased TGFβ1-induced migration and lung colonization upon MR knockdown parallels the reduced migration, invasion, and lung colonization we observed when GR Ser134 was mutated to alanine (S134A GR models). This led to the formulation of our hypothesis that MR and p-GR work in cooperation under the influence of growth factors and cytokines abundantly present in the tumor-microenvironment. We tested this hypothesis via the proximity ligation assay, which is a powerful tool to detect in situ protein-protein interactions (84). Recent studies using proximity ligation assay (PLA) have provided direct evidence for MR-GR interactions at the cellular level. For instance, PLA was employed to demonstrate MR-GR interactions in hippocampal neurons under conditions of corticosterone stimulation, revealing a mechanism by which stress hormones can fine-tune receptor signaling (85). Similarly, PLA was used to detect MR-GR complexes in brain tissue, showing that receptor interaction patterns change in response to stress and contribute to synaptic plasticity (86). Through NanoBiT assays in U2OS (osteosarcoma) cell line models where MR and GR are overexpressed, it was shown that with a simultaneous addition of dex and aldo MR-GR interaction was significantly enhanced compared to the single ligand effects (87). These examples align with our results from TNBC cell lines, in which we also observed increased MR-GR complex formation upon combined dex and aldo treatment. Moreover, both spiro and fine reduced MR-GR interaction puncta that are induced by single hormone ligands or the combination of both. These results are consistent with data in multiple myeloma cells that dex mediated MR-GR interactions are reduced from 8-fold to 6-fold upon spiro treatment (45). With TGFβ1 treatment we observed that the complexes were initially more cytoplasmic than nuclear (90 min), but the MR-GR co-complexes translocated to the nucleus within 4 hours. We speculate that these complexes may interface with rapidly activated cytoplasmic signaling pathways organized by 14-3-3ζ, which appears to serve a scaffolding function that nucleates and/or stabilizes functional p38 MAPK modules, thereby enabling ultrasensitive signal transduction in response to a wide range of cellular stress stimuli (88) (Figure 7C).

Our studies have revealed two distinct regulatory MR-GR axes downstream of aldo, dex and TGFβ1: (1) a p38 MAPK–dependent, GR Ser134 phosphorylation-dependent step required for MR-p-GR binding in the cytoplasm that ultimately becomes nuclear, and (2) a more rapid ligand-dependent MR nuclear import that can both be blocked by MR antagonists. This dual regulation of MR signaling induced by either cytokines or steroid hormones highlights that cytoplasmic MR-p-GR association under profibrotic or other inflammatory cues is not merely a byproduct of MR nuclear shuttling, but rather an independent active, kinase-driven event that may have the same/similar or totally distinct functional consequences. Likewise, both aldo and TGFβ1 induced tumorsphere formation and cell migration through MR-dependent mechanisms; however, the nature and localization of MR-GR complexes differed between the stimuli. Hormone treatments primarily promoted the formation of nuclear localized complexes, whereas cytokine stimulation resulted in cytoplasmic associations. Interestingly, MAP3K5, an upstream activator of the p38 MAPK signaling cascade, was significantly upregulated at the mRNA level following 24 hours of TGFβ1 treatment, but this induction was markedly reduced upon MR knockdown. These findings suggest that MR contributes to TGFβ1-mediated activation of MAP3K5 and downstream p38 signaling, potentially linking MR-dependent cytoplasmic signaling to cytokine-induced migratory phenotypes while driving a feed-forward TGFβ signaling loop that sustains stress signaling. Given that p38 MAPK activation has been associated with epithelial–mesenchymal transition (EMT), cytoskeletal remodeling, and tumor cell invasiveness, these findings suggest that MR may serve as a key mediator integrating cytokine cues into pro-migratory and stress-response signaling networks (88).

Thus, we propose that while nuclear MR-GR complexes may mediate transcriptional responses to hormones, the cytoplasmic MR-GR complexes formed in response to cytokines, such as TGFβ1, may promote non-genomic signaling through MAP3K5–p38 MAPK, with each mechanism contributing to aggressive tumor phenotypes, perhaps via common signaling or transcriptional convergence points. Despite the differences in initial subcellular localization patterns, both aldo- and TGFβ1-induced complexes promoted tumorsphere formation and enhanced migratory capacity in an MR-dependent manner. These data indicate that MR serves as a central integrator of both endocrine and inflammatory cues, activating distinct signaling routes. Under hormone-rich conditions, the nuclear co-complexes of MR-GR exert rapid transcriptional responses, whereas the cytokine-dependent, cytoplasmic, non-genomic responses activate downstream cell signaling events that ultimately drive similar pro-tumorigenic phenotypes. Biologically, this dual mode of MR signaling provides a mechanistic framework for how stress hormones and inflammatory cytokines cooperate to promote cancer cell migration and tumorsphere formation under pathophysiological conditions. Studies of global MR-pS134-GR-dependent vs. MR-GR DNA-binding and gene expression are the topic of ongoing future studies. In summary, our novel study highlights that MR activation by either its physiological ligand aldosterone or by cytokines (i.e., TGFβ1) leads to increased migration and stem-cell like phenotypes in TNBC models that are efficiently blocked by FDA-approved MR antagonists. As such, MR contributes to efficient TNBC lung colonization. These findings offer new insights into the potential therapeutic possibilities and feasibility for targeting the MR-p-GR axis using selective MRAs (finerenone) in the near-term to block invasive and metastatic tumor behaviors in patients with TNBC.

## Author Contributions

S.H.P., C.H.D., R.I.K., and H.S. performed the experiments. S.H.P. and C.H.D. analyzed and interpreted the data with input from T.N.S, J.B., and C.A.L. S.H.P., C.H.D., T.N.S., J.B., and C.A.L. conceptualized the study. C.A.L. conceived the study. S.H.P., C.H.D., T.N.S., and C.A.L. wrote and/or edited the manuscript. C.A.L. supervised the study. All authors read and reviewed the final manuscript and consented to publication of the manuscript.

## Abbreviations

Aldo: aldosterone
ANOVA: analysis of variance
BC: breast cancer
BSA: bovine serum albumin
cDNA: complementary DNA
DCC: steroid hormone-free fetal bovine serum
Dex: dexamethasone
DMEM: dulbecco’s modified eagle’s medium
DMSO: dimethyl sulfoxide
EGF: epidermal growth factor
EMT: epithelial to mesenchymal transition
ER+: estrogen receptor positive
FBS: fetal bovine serum
Fine: finerenone
GR: glucocorticoid receptor
HER2: human epidermal growth factor
IMEM: improved minimal essential medium
MAPK: mitogen-activated protein kinase
MEM: minimal essential medium
METABRIC: molecular taxonomy of breast cancer international consortium
MR: mineralocorticoid receptor
mRNA: messenger RNA
PAGE: polyacrylamide gel electrophoresis
PBS: phosphate-buffered saline
PLA: proximity ligation assay
PR: progesterone receptor
pS134-GR: phospho-Ser134 glucocorticoid receptor
qPCR: quantitative polymerase chain reaction
SD: standard deviation
SDS-PAGE: sodium dodecyl sulfate-PAGE
SR: steroid hormone receptor
TGFβ1: transforming growth factor β1
TME: tumor microenvironment
TNBC: triple-negative breast cancer
WT: wild-type

