## Supplementary Figures for "Mineralocorticoid and glucocorticoid receptor interaction drives TGFβ1-induced triple-negative breast cancer progression to metastasis"

### Supplementary Figure 1

A. Primer sequences for RT-qPCR analysis

| | | |
|---|---|---|
| MR | F | CACCATGCAGGCAACATTAC |
| | R | CTGAGTGCAGCTTTCACAGG |
| SGK1 | F | TCTCAGCAAATCAACCTTGG |
| | R | TCTTCTGCCTTGTGTCTTGC |
| ENAC1 | F | TGCACCTGTCAGGGGAAC |
| | R | CTTCATGAGCCCTGGAGTGG |
| COL1A1 | F | TCAGGGAATGCCTGGTGAAC |
| | R | GGACCAGCATCACCTCTGTC |
| ICAM1 | F | AGCTTCGTGTCCTGTATGGC |
| | R | GGAATTTTCTGGCCACGTCC |
| CTGF | F | GGGCCTATTCTGTCACTTCGG |
| | R | AGCACCATCTTTGGCGGTG |
| EDN1 | F | CTACTTCTGCCACCTGGACATC |
| | R | TCACGGTCTGTTGCCTTTGTGG |
| CLDN14 | F | CTGGACTTGGCTGAGGACAC |
| | R | GACTGCGGCTTCAACAGTTC |
| MAP3K5 | F | AGGTGGTACTCTTTGGTTTTCAAG |
| | R | GATACTGTCTAAGGCAAACATCCAG |
| 18 | F | GGAGAGGGAGCCTGAGAAAC |
| | R | TCGGGAGTGGGTAATTTGC |

**Supplementary Figure 1**
(A) Primer sequences used for RT-qPCR analysis

### Supplementary Figure 2

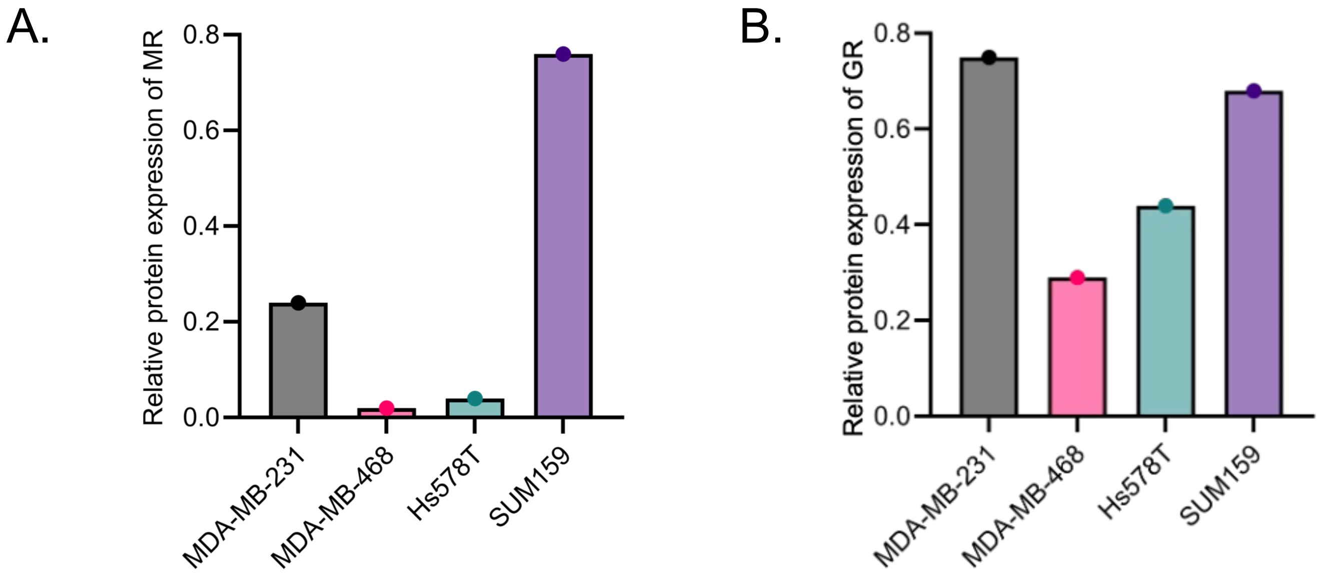


**<u>Supplementary Figure 2</u>**

(A) Densitometry values of relative protein expression of MR in Triple-Negative breast cancer (TNBC) cell lines shown in Figure 1A. (B) Densitometry values of relative protein expression of GR in TNBC cell lines shown in Figure 1A.

### Supplementary Figure 3

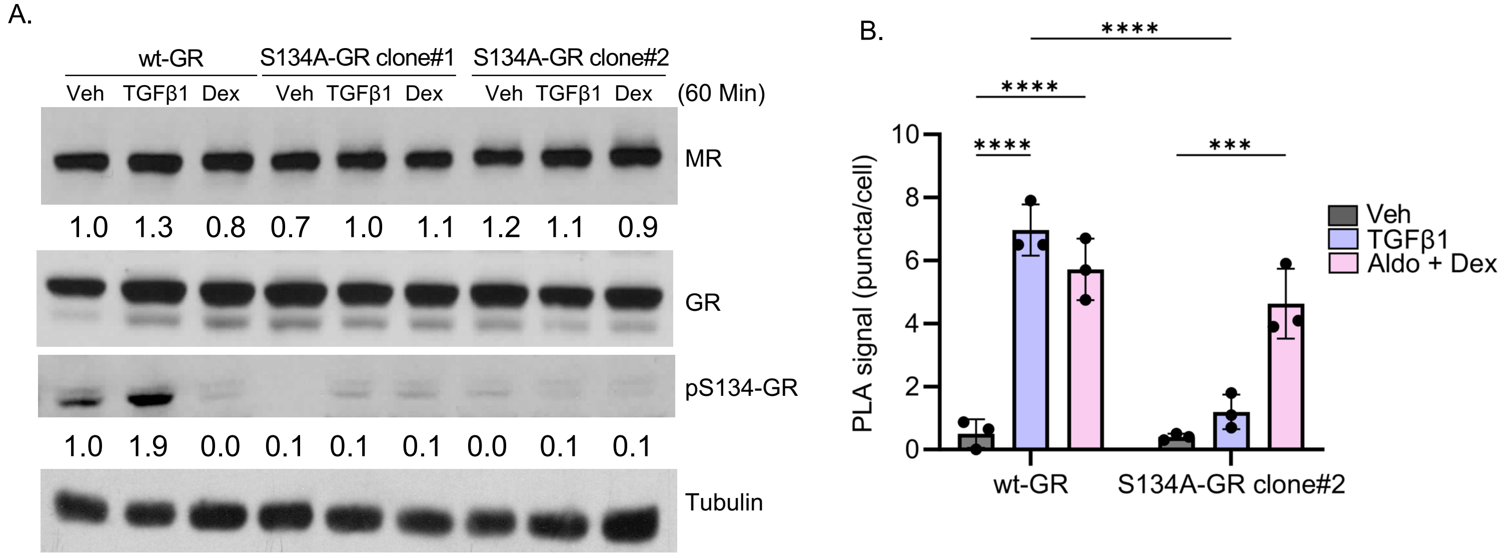


**<u>Supplementary Figure 3</u>**

(A) Protein expression of MR, GR, pS134-GR and tubulin in wt-GR, S134A-GR clone #1 and clone #2 treated with 10ng/ml TGFβ1 or 1 uM dexamethasone (dex) for 60 min. (B) Quantification of the red PLA puncta showing an interaction between MR and GR in wt-GR and S134A-GR clone #2 MDA-MB-231 cells treated with 10ng/ml TGFβ1 or 1uM aldosterone (aldo) plus 1 uM dexamethasone (dex) for 90 min and Error bars are S.E.M.; n = 3 biological replicates; **P < 0.01; ***P < 0.001; ****P < 0.0001.

### Supplementary Figure 4

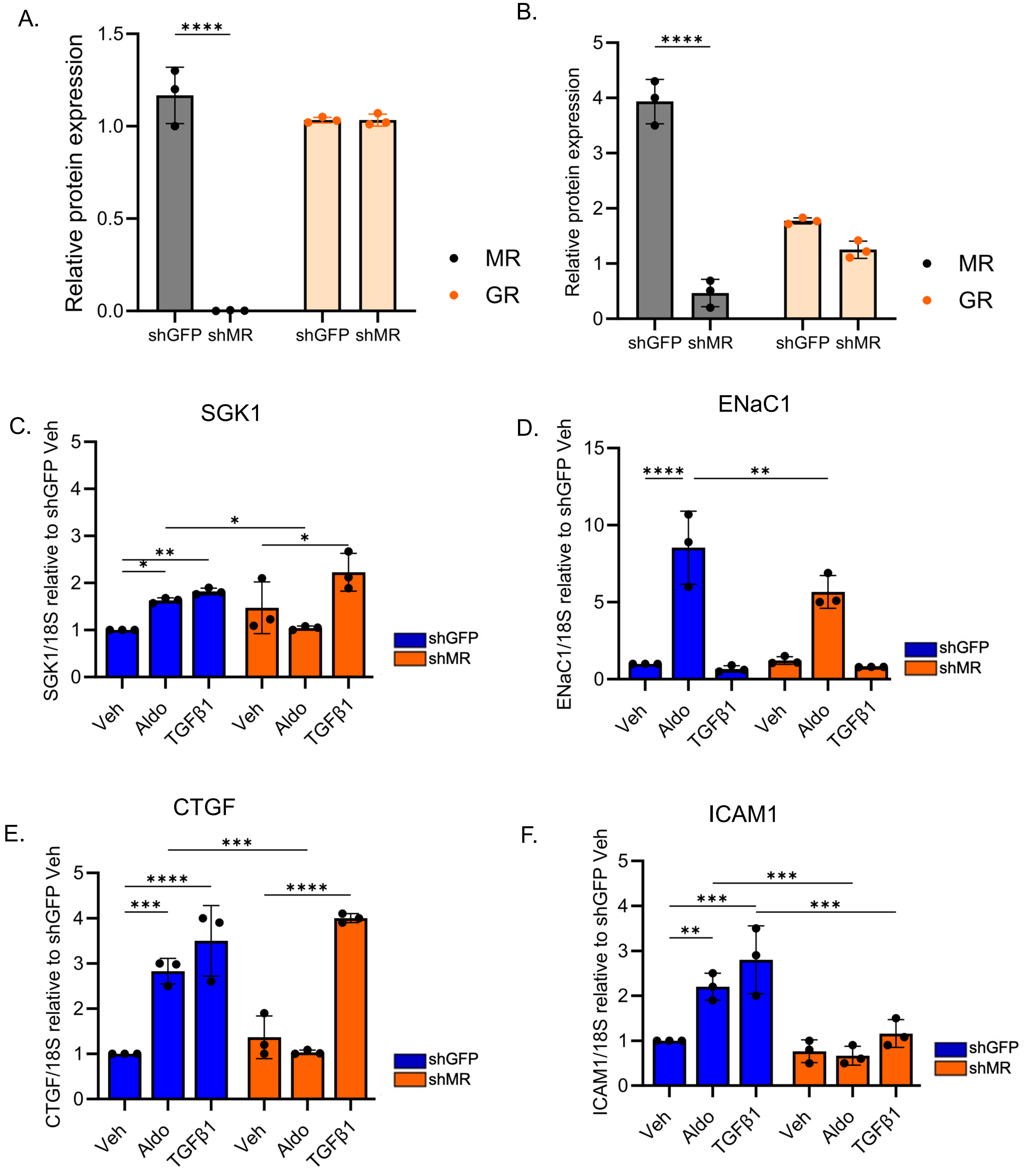


**Supplementary Figure 4**

(A-B) Densitometry analysis of relative protein expression of MR and GR in shGFP and shMR (A) MDA-MB-231 OR (B) SUM159 (B) cells. (C-F) mRNA levels of (C) SGK1, (D) ENaC1, (E) CTGF, and (F) ICAM1 assessed by RT-qPCR in shGFP and shMR SUM159 cells treated with 1 µM aldosterone (aldo) or 10 ng/ml TGFβ1 for 6 hours. The mean of three biological replicates is shown ± SD. (*, P < 0.05, **, P < 0.01, ***, P < 0.001, ****, P < 0.0001).

### Supplementary Figure 5

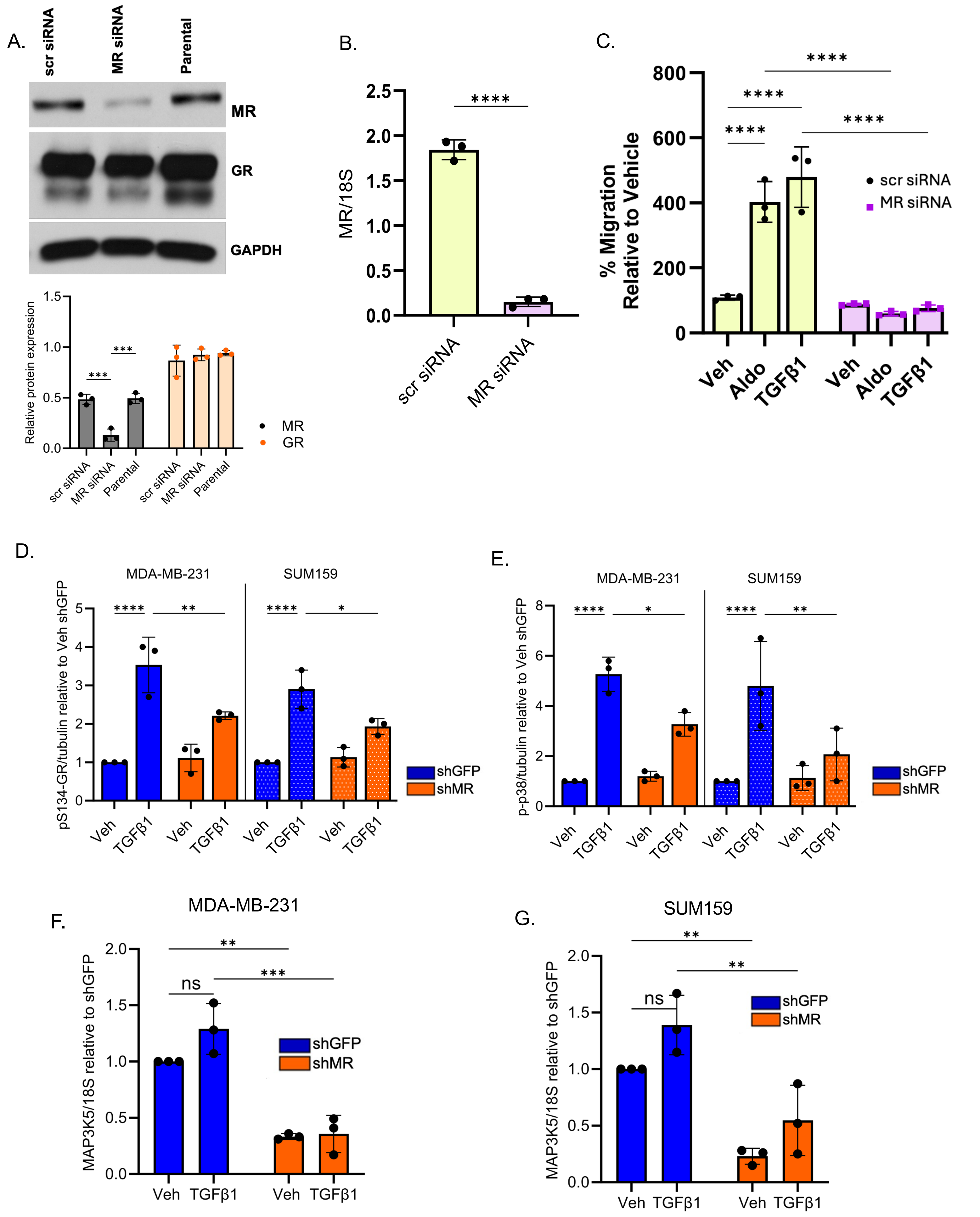

**Supplementary Figure 5**

(A) Western blot analysis of MR and GR levels in MDA-MB-231 cells transfected with either scramble (scr) siRNA or MR siRNA compared to the parental cells and densitometry values. (B) MR mRNA levels in MDA-MB-231 cells exposed scramble (scr) siRNA or MR siRNA were measured by RT-qPCR following normalization to 18S mRNA. Mean expression of three independent replicates ± SD is shown. (C) Transwell migration assays of MDA-MB-231 scr siRNA or MR siRNA cells treated with either vehicle (veh), aldosterone (aldo) or TGFβ1 (10 ng/mL) for 18 hours using 10% FBS as a chemoattractant. (D-E) Densitometry analysis for (D) pSer134-GR or (E) p-p38 normalized to total GR or total p38 for the western blot shown in Figure 6. (F-G) mRNA levels of MAP3K5 were assessed by RT-qPCR in shGFP and shMR (F) MDA-MB-231 or (G) SUM159 cells treated with 10 ng/ml TGFβ1 for 6 hours, following normalization to 18S mRNA and to the normalized vehicle control (set to 1.0). The Mean of three biological replicates is shown ± SD: *, $P < 0.05$, **, $P < 0.01$, ***, $P < 0.001$, ****, $P < 0.0001$.

### Supplementary Figure 6

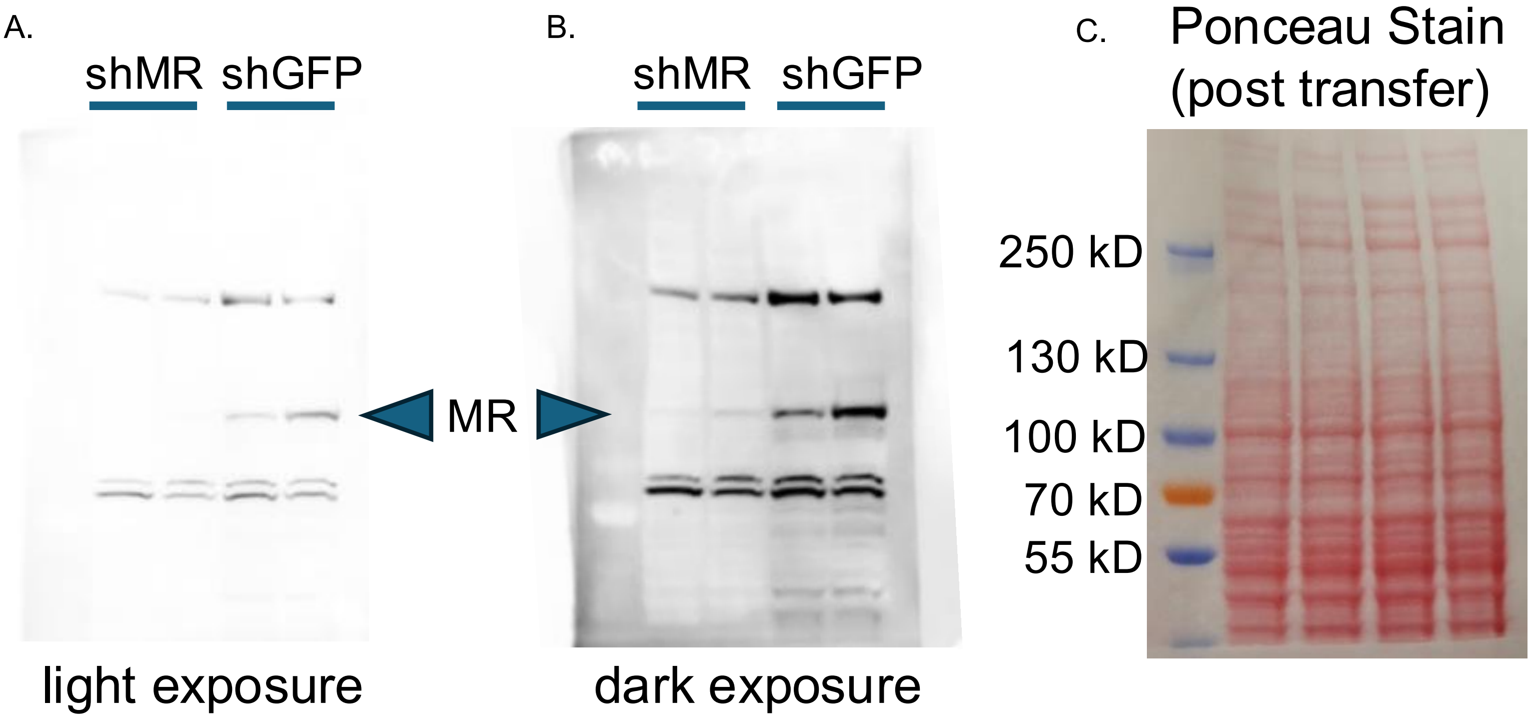


**Supplementary Figure 6**

Western blot analysis of MR protein expression in MDA-MB-231 shGFP and shMR cells used for mouse tail vein injection experiments. (A) Short exposure and (B) long exposure of MR expression in the Western blot analysis. (C) Ponceau S staining of Western blot membrane.
